# Early activation of cellular stress and death pathways caused by cytoplasmic TDP-43 in the rNLS8 mouse model of ALS/FTD

**DOI:** 10.1101/2022.08.08.503119

**Authors:** Wei Luan, Amanda L. Wright, Heledd Brown-Wright, Sheng Le, Rebecca San Gil, Lidia Madrid San Martin, Karen Ling, Paymaan Jafar-Nejad, Frank Rigo, Adam K. Walker

## Abstract

TAR DNA binding protein 43 (TDP-43) pathology is a key feature of over 95% of amyotrophic lateral sclerosis (ALS) and nearly half of frontotemporal dementia (FTD) cases. The pathogenic mechanisms of TDP-43 dysfunction are poorly understood, however activation of cell stress pathways may contribute to pathogenesis. We therefore sought to identify which cell stress components are critical for driving disease onset and neurodegeneration in ALS/FTD. We studied the rNLS8 transgenic mouse model, which expresses human TDP-43 with a genetically-ablated nuclear localisation sequence within neurons of the brain and spinal cord resulting in cytoplasmic TDP-43 pathology and progressive motor dysfunction. Amongst numerous cell stress-related biological pathways profiled using qPCR arrays, several critical ISR effectors, including CCAAT/enhancer-binding homologous protein (*Chop/Ddit3*) and activating transcription factor 4 (*Atf4*), were upregulated in the cortex of rNLS8 mice prior to disease onset. This was accompanied by early up-regulation of anti-apoptotic gene *Bcl2* and diverse pro-apoptotic genes including BH3-interacting domain death agonist (*Bid*). However, pro-apoptotic signalling predominated after onset of motor phenotypes. Notably, pro-apoptotic caspase-3 protein was elevated in the cortex of rNLS8 mice at later disease stages, suggesting that downstream activation of apoptosis drives neurodegeneration following failure of early protective responses. Unexpectedly, suppression of *Chop* in the brain and spinal cord using antisense oligonucleotide-mediated silencing had no effect on overall TDP-43 pathology or disease phenotypes in rNLS8 mice. Cytoplasmic TDP-43 accumulation therefore causes very early activation of ISR and both anti-and pro-apoptotic signalling that switches to predominant pro-apoptotic activation later in disease. These findings suggest that precise temporal modulation of cell stress and death pathways may be beneficial to protect against neurodegeneration in ALS and FTD.

**Key points:**

1. ISR genes *Atf4* and *Chop,* anti-apoptotic Bcl2 and pro-apoptotic gene *Bid*, *Bim*, *Noxa* were upregulated in the cortex of rNLS8 mice prior to disease onset
2. Knockdown of *Chop* had limited effects on TDP-43 pathology and did not alter motor deficits in rNLS8 mice
3. Both anti-and pro-apoptotic genes are upregulated prior to disease onset, and switches to activation of pro-apoptotic signalling at later disease stages
4. Caspase-3 activation likely drives neurodegeneration in the cortex of rNLS8 mice

## Introduction

Amyotrophic lateral sclerosis (ALS) and frontotemporal dementia (FTD) are neurodegenerative diseases that share multiple pathological features, particularly the presence of intracellular protein inclusions. Of proteins contributing to disease, ubiquitinated, phosphorylated, and biochemically-insoluble forms of TAR DNA binding protein 43 (TDP-43) lead to neurodegeneration in >95% of both sporadic and familial ALS cases and about 45% of FTD cases (1, 2). TDP-43 is an important RNA-binding protein that typically shuttles between the nucleus and cytoplasm under physiological conditions, but TDP-43 accumulation in the cytoplasm of neurons results in neurodegeneration (3, 4). The pathways leading to neuronal demise caused by cytoplasmic TDP-43 pathology remain to be fully defined (5), and understanding the mechanisms that cause neuron death may reveal new therapeutics opportunities.

Many pathogenic mechanisms have been proposed to contribute to neurodegeneration in ALS, including the integrated stress response (ISR) (6), protein synthesis dysregulation (7), oxidative stress (6), and neuroinflammation (8, 9). Indeed, chronic activation of the ISR likely contributes to motor neuron death in ALS/FTD (10, 11). The ISR is a master regulatory pathway that fine-tunes proteostasis in cells, controlling protein synthesis, folding, and decay processes, and is crucial for cell survival (11). Under diverse cellular stress conditions, ISR signaling upregulates select downstream transcripts to restore protein homeostasis. Activating transcription factor 4 (ATF4) is a crucial ISR regulator that targets genes to promote protein folding and degradation, increasing cell survival (12). Nevertheless, prolonged disturbance of proteostasis enhances ISR signaling and consequently results in ATF4-mediated upregulation of CCAAT/enhancer-binding homologous protein (CHOP), which can trigger a pro-apoptotic signal cascade (11, 13). ISR activation, evidenced by activation of ISR sensor kinases, has been reported in postmortem samples of ALS cases (6). Moreover, activation of ISR enhances TDP-43 aggregation and stress granule formation, which are hallmarks of TDP-43-linked disease (14–16), suggesting a direct association of TDP-43 dysfunction with ISR activation. In addition, TDP-43 toxicity increases levels of *CHOP protein* in cell lines and sporadic ALS spinal cord tissues(15, 17). Pharmacological modulation for ISR using small-molecule drugs can modify neurodegeneration (7), however, the actions of these drugs are not restricted to the ISR pathways. Therefore, it remains unclear whether specific modulation of ISR effectors can modulate disease in mammalian models of cytoplasmic TDP-43 proteinopathy.

Transgenic mice with doxycycline (Dox)-suppressible expression of human TDP-43 (hTDP-43) containing a defective nuclear localization signal under the control of the neurofilament heavy chain promoter (rNLS8 mice) are one of the most disease relevant models of ALS, displaying progressive TDP-43 cytoplasmic accumulation coinciding with rapid ALS-like neurodegeneration and motor decline [15, 16]. In this study, we therefore screened for genes that may regulate and respond to cellular stress pathways in the cortex of rNLS8 mice prior to disease onset and throughout early disease stages. We thereby identified alterations in expression of several genes across multiple molecular pathways, including upregulation of genes involved in the ISR, DNA damage response, apoptosis and neuroinflammation, and downregulation of genes involved in glycolysis and ion exchange, in rNLS8 mice. Remarkably, several genes involved in the ISR, DNA damage response and apoptosis were dramatically increased even prior to disease onset. Notably, anti-and pro-apoptotic genes were up-regulated prior to disease onset, with a switch towards solely pro-apoptotic signaling at later disease stages in rNLS8 mice. Our results also revealed a significant elevation of caspase-3 levels in late-disease rNLS8 mice, supporting ISR-mediated apoptosis as a critical contributor to neurodegeneration in disease. To investigate whether amelioration of ISR signaling could protect against disease, we therefore performed intracerebroventricular (ICV) injection of antisense oligonucleotides (ASOs) to suppress *Chop* expression (18), in rNLS8 mice. ASO treatment decreased *Chop* levels but did not affect overall TDP-43 pathology, neurodegeneration, gliosis, or motor deficits in rNLS8 mice. Taken together, we conclude that TDP-43 mislocalisation leads to early perturbation of multiple molecular pathways that crosstalk with ISR signals, with dysregulation of apoptosis signaling contributing to disease in rNLS8 mice.

## Results

### Early activation of cell stress and death pathways in the cortex of rNLS8 mice

To determine how numerous cell stress and death pathways are regulated in the cortex of rNLS8 mice in early disease stages, we conducted RT^2^ PCR arrays to assess the expression of critical genes at time of approximate disease onset (2 weeks off Dox, 2 WOD) and an early disease stage (4 weeks off Dox, 4WOD) (Fig 1A). We assessed 84 mouse genes in ten functional groups, with four housekeeping genes *(B2m* excluded) and three negative controls (the full gene expression dataset is available in Supplementary Table 1). Firstly, we confirmed dramatic increases of the human *TARDBP (hTARDBP)* mRNA levels in disease rNLS8 mice at both 2 and 4 WOD relative to control mice (Fig 1B) using real-time qPCR, as expected. In contrast, the mRNA levels of mouse *TARDBP (mTARDBP)* were not significantly different between control and rNLS8 mice in this bulk tissue analysis (Fig 1B), although loss of endogenous TDP-43 protein from neurons is also a disease-reminiscent feature of this model (19).

**Fig. 1.**
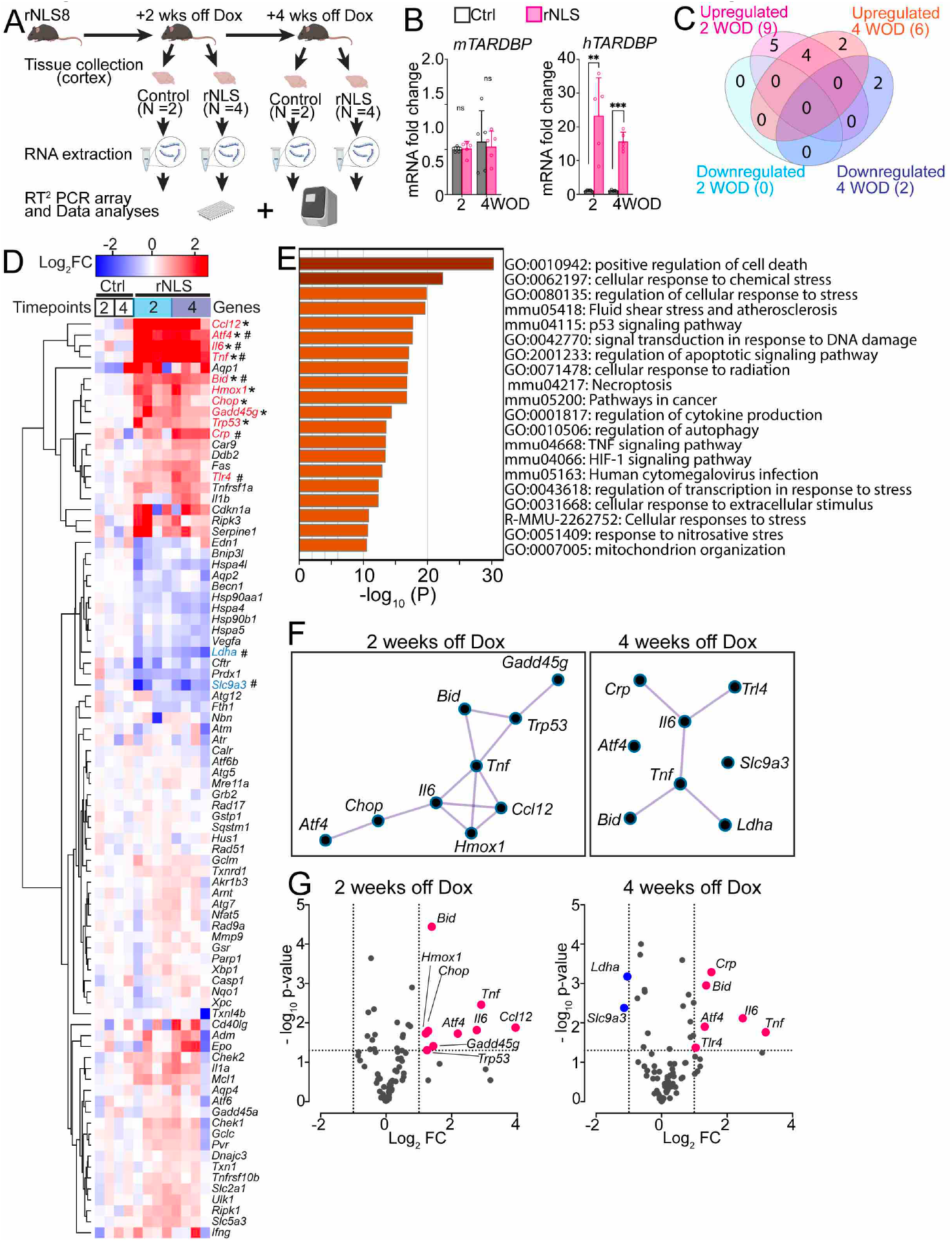
Cellular stress and death signaling pathways are activated in the cortex of rNLS8 mice at early disease stages. **A**. Experimental schema. **B**. Real-time qPCR analysis of mouse and human TARDBP gene expression in the cortex of rNLS8 mice at 2 and 4 weeks off Dox (WOD) relative to controls. n = 5. Mean + SEM. **p < 0.01 and ***p < 0.001. **C**. Total number of unique and shared statistically significantly upregulated or downregulated genes in rNLS8 mice at each time point, relative to the control mice. **D.** Unsupervised hierarchical clustering heatmap for expression of all assayed genes in RT^2^ PCR array from rNLS8 and control mice at each timepoint, with values given as normalised gene expression levels (log2 fold changes). Statistically significantly (p < 0.05) upregulated and downregulated genes are indicated in red and blue, respectively. * indicates statistical significance for 2 WOD and # for 4 WOD. **E.** Gene ontology enrichment analyses for all the genes of interest in in RT^2^ PCR arrays. **F.** MCODE network analyses for protein-protein interaction of identified differentially expressed genes in rNLS8 mice at 2 and 4 WOD, respectively. **G.** Volcano plots indicate the magnitude (x axis, as log_2_ fold changes) and statistical significance (y axis, as-log_10_ p-values) of gene expression changes in rNLS8 mice relative to control mice at each timepoint. Upregulated genes (fold change > 2) are shown in red and downregulated genes (fold change < −2) are shown in blue.

**Table 1.**
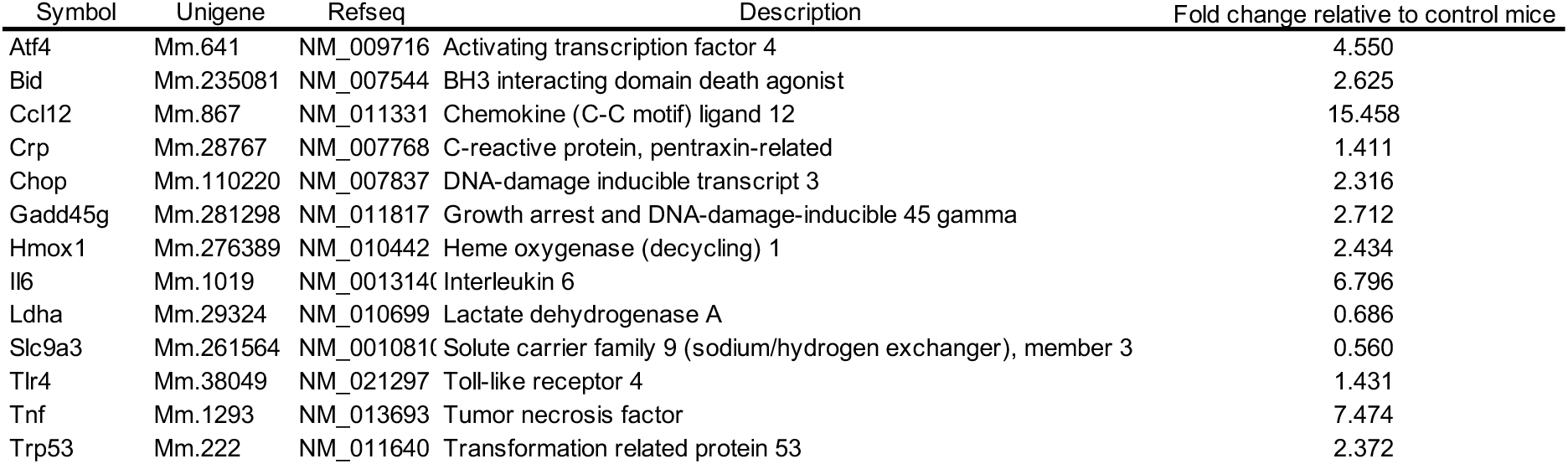
Genes showing statistical significant difference in cortex of rNLS8 mice at 2 weeks and/or 4 weeks off Dox compared to controls

**Table 2.**
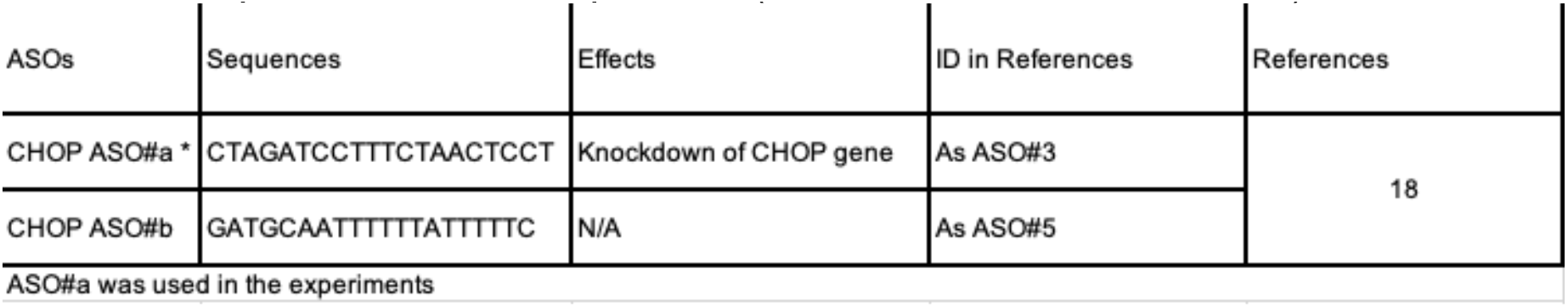
Sequences of the *Chop* ASOs (available in Excel document)

At 2 weeks off Dox, taken as disease onset, we identified a total of nine statistically upregulated genes (fold change > 2, *p* < 0.05) in the rNLS8 mice (Fig 1C, the expression dataset of identified genes is available in Table 1). At 4 weeks off Dox, taken as an early disease stage after accumulation of TDP-43 pathology, we detected six upregulated genes and two downregulated genes (fold change < −2, p < 0.05) in the rNLS8 mice relative to controls (Fig 1C). Among these dysregulated genes, we identified four genes that were upregulated at both 2-week and 4-week timepoints in the cortex of rNLS8 mice in the RT2 array, namely activating transcription factor 4 *(Atf4),* pro-apoptotic BH3 Interacting Domain Death Agonist *(Bid),* interleukin 6 *(Il6)* and tumor necrosis factor-alpha *(Tnf)*.

The hierarchical heatmap showing the magnitude of fold change in expression between rNLS8 mice and controls in the RT^2^ array demonstrated a clear bi-clustering pattern of upregulated genes (in red) and downregulated genes (in blue) over the disease course (Fig 1D). The GO enrichment analyses of all pre-selected genes revealed that the top three biological processes relate to ISR signaling: positive regulation of cell death (GO:0010942), cellular response to chemical stress (GO:0062197), and regulation of cellular response to stress (GO:0080135) (Fig 1E). Furthermore, the molecule complex detection (MCODE) network for protein-protein interaction revealed significant interactions of identified genes among biological processes, notably the positive regulation of cell death via the ATF4-CHOP axis of the ISR in the disease-onset rNLS8 mice at 2 weeks off Dox (Fig 1F).

To further interrogate the gene changes in rNLS8 mice, we grouped analysed genes based on biological pathway at both 2-week and 4-week timepoints (Figure 2). At 2 weeks off Dox, upregulated genes act within several pathways: the ISR, being *Atf4* (~3.7 fold) and C/EBP homologous protein *(Chop*, ~2.3 fold) (Fig 2A); apoptosis signaling, being *Bid* (~2.6 fold) (Fig 2B); oxidative stress response, being Heme Oxygenase 1 (*Homx1*, ~2.4 fold) (Fig 2C); DNA damage response, being DNA Damage Inducible Gamma *(Gadd45g*, ~2.7 fold) and transformation related protein 53 (*Trp53*, ~2.3 fold) (Fig 2D); neuroinflammation, being chemokine (CC motif) ligand 12 (*Ccl12*, ~15.5 fold) and *Il6* (~6.7 fold) (Fig 2E), and; TNF family signaling, being *Tnf* (~7.4 fold) (Fig 2F). The data suggest that TDP-43 cytoplasmic mislocalisation in neurons of rNLS8 mice results in disruption of cellular homeostasis including ISR activation, DNA damage, oxidative stress, and neuroinflammation as early as disease onset.

**Fig. 2.**
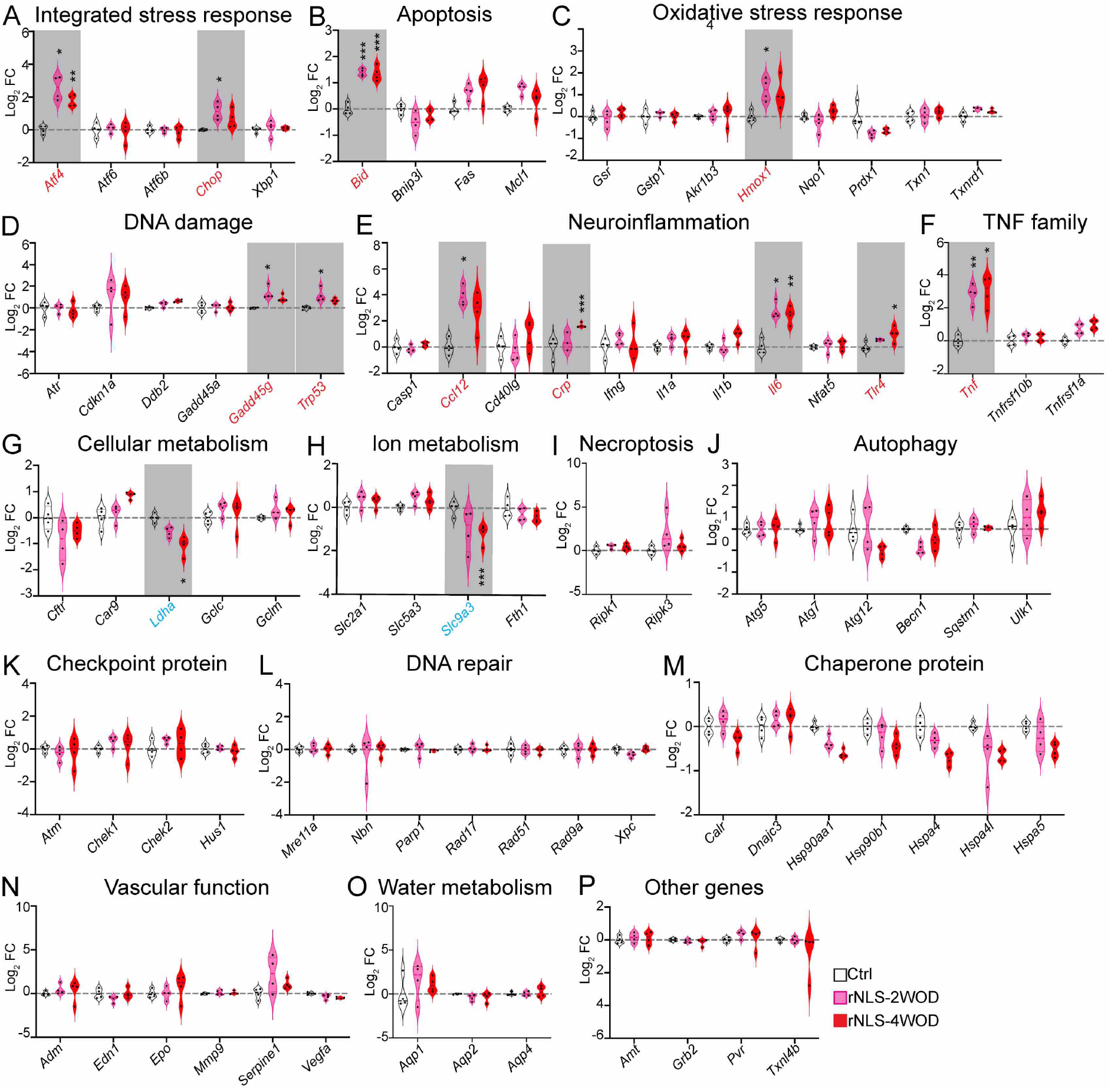
rNLS8 mice display significant dysregulation of genes in multiple cellular stress pathways in the cortex at early disease stages. The expression profiles for individual assayed genes from RT^2^ PCR array analyses assigned to different biological functional groups at 2 and 4 weeks off Dox (WOD). **A**. Integrated stress response pathway. **B**. Apoptosis signaling. **C**. Oxidative stress response. **D**. DNA damage response. **E**. Neuroinflammation signaling. **F**. TNF family signaling. **G**. Cellular metabolism. **H**. Ion metabolism. **I**. Necroptosis. **J**. Autophagy. **K**. Checkpoint protein. **L**. DNA repair. **M**. Chaperone protein. **N**. Vascular function. **O**. Water metabolism. **P**. Remaining other genes. The statistically upregulated genes (labels in red) and the downregulated genes (in blue) are highlighted in grey boxes. n = 4. *p < 0.05, **p < 0.01 and ***p < 0.001.

At 4 weeks off Dox, upregulated genes largely belong to the same biological pathways as at 2 weeks: the ISR, being *Atf4* (~ 2.5 fold) (Fig 2A); apoptosis signaling, being *Bid* (~2.5 fold) (Fig 2B); neuroinflammation, being the C-reactive protein gene (*Crp*, ~2.8 fold), Il6 (~ 5.5 fold), and the Toll-like receptor 4 gene *(Tlr4,* ~2.1 fold) (Fig 2E), and; TNF family ligand, being *Tnf* (~ 9.0 fold) (Fig 2F). Furthermore, we also identified two downregulated genes in in the cortex of rNLS8 mice at 4 weeks off Dox that were not altered at 2 weeks, involved in: cellular metabolism (glycolysis), being lactate dehydrogenase A (*Ldha*, ~ 0.5 fold) (Fig 2 G), and; ion metabolism, being the solute carrier family 9 member A3 (*Slc9a3*, ~0.5 fold) (Fig 2H). These results suggest there is a prolonged elevation of cell stress response during disease and later dysregulation in cellular metabolism in the rNLS8 mice.

Notably, four genes that were upregulated at both 2-week and 4-week timepoints in the cortex of rNLS8 mice in the RT^2^ array are the ISR gene *Atf4* (Fig 2A), the apoptotic gene *Bid* (Fig 2B), neuroinflammatory gene *Il6* (Fig 2E), and TNF family gene *Tnf* (Fig 2F). Consistent alteration in these pathways indicates involvement of these genes in disease over an extended period, suggesting that these pathways may be important during disease progression.

To validate our findings in RT^2^ array analyses and determine whether changes occur even before disease onset, we conducted real-time qPCR for all 13 genes detected as significantly altered in at least one of the timepoints in the rNLS8 mice, in separate experiments using an independent set of mouse cortex samples. In addition to disease onset (2 weeks) and early disease (4 weeks) timepoints, we additionally analysed samples from mice prior to disease onset (1 week off Dox), a timepoint at which rNLS8 mice display increased cytoplasmic human TDP-43 protein levels but no overt motor phenotypes (19). Intriguingly, five of the 11 upregulated genes originally identified as altered in array data were already upregulated at 1 week timepoint (Fig 3A), including genes involved in: the ISR, being *Atf4* (~ 1.5 fold) and *Chop* (~ 2.1 fold); apoptosis signaling, being *Bid* (~ 2.2 fold), and; DNA damage response, being *Gadd45g* (~ 1.4 fold) and *Trp53* (~ 1.7 fold). Expression of these five genes were also consistently elevated at 2 weeks and 4 weeks off Dox relative to the control mice, consistent with the array findings. Consistent upregulation of these genes suggests the early and prolonged activation of cellular stress pathways and pro-apoptosis signaling in the cortex of rNLS8 mice upon the induction of *hTDP-43^ΔNLS^* expression, even before disease onset.

**Fig. 3.**
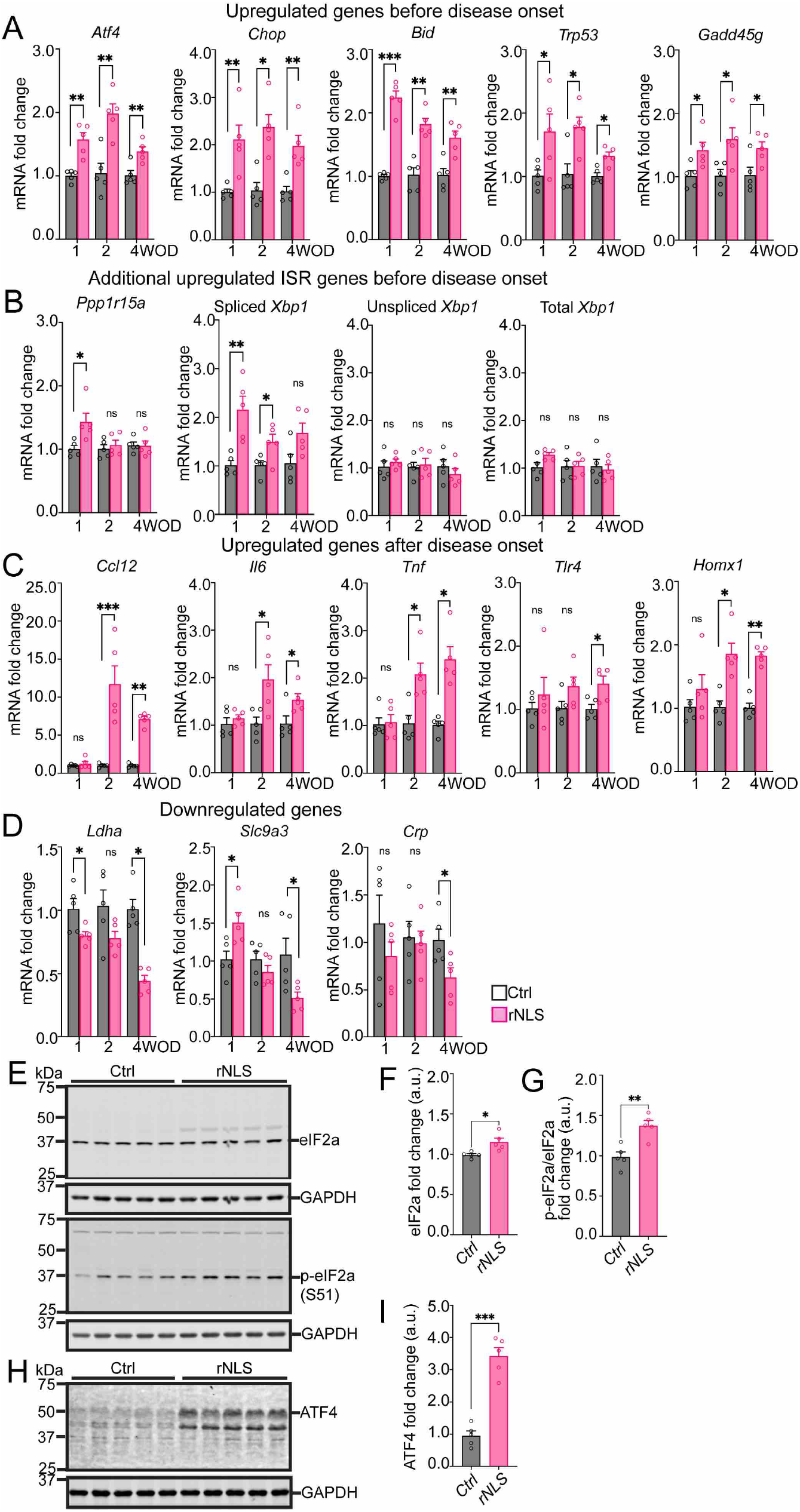
rNLS8 mice display dysregulation of genes in multiple cellular stress pathways and activation of the ISR in the cortex even prior to disease onset. (**A**) Real-time qPCR verified upregulated RT^2^ array-identified genes (Atf4, Chop, Bid, Trp53 and Gadd45g) in the cortex of rNLS8 mice at 2 and 4 weeks off Dox (WOD), and revealed similar upregulation at 1 week off Dox. (**B)** Additional ISR-related genes (Ppp1r15a and spliced Xbp1) were also upregulated at 1 WOD. (**C**) The RT^2^ array-identified upregulated genes Ccl12, Il6, Tnf, Tlr4 and Homx1 were verified as increased at 2 and 4 WOD but were unaltered at 1 WOD. (**D**) Differential alterations were also identified in the RT^2^ array-identified downregulated genes (Lhda, Slc9a3 and Crp) at 1 and 4 WOD. Values were normalized to Actb as the housekeeping gene. **E**. Immunoblotting for the proteins of eIF2α and p-eIF2α (S51) in in the cortex of rNLS8 mice at 4 WOD. Approximate molecular weights (kDa) are indicated. Quantification of immunoblots showed increased eIF2α (**F**) and p-eIF2a (**G**) in the cortex of rNLS8 mice relative to control mice, respectively. (**H)** Immunoblotting for ATF4 protein in the cortex of rNLS8 mice at 4 WOD. (**I)** Quantification of immunoblots showed increased ATF4 protein in the cortex of rNLS8 mice relative to control mice. n = 5. Mean + SEM. *p < 0.05, **p < 0.01, ***p < 0.001.

Given the prominence of changes in the ISR pathway early in the rNLS8 mice, we also examined changes in additional ISR genes not included in the array data, including levels of protein phosphatase 1 regulatory subunit 15A mRNA *(Ppp1r15a)* that is the direct target of ISR-ATF4-CHOP axis (11), and mRNA splicing of X-box binding protein 1 *(Xbp1)* that is a potent transcription factor under regulation of the inositol-requiring enzyme 1 (IRE1) branch of unfolded protein response (UPR) (12). Notably, we identified upregulation of the *Ppp1r15a* gene encoding GADD34 (~ 1.4 fold, Fig 3B) at 1-week timepoint, but not 2-or 4-week timepoint, suggesting impairment of GADD34-mediated negative feedback regulation for the ISR to recovering from translational inhibition (20). As expected, the total and unspliced *Xbp1* mRNAs showed no significant differences between groups (Fig 3B), in line with the RT^2^ array data for total *Xbp1* mRNAs (Fig 2A). However, there was marked elevation of spliced (active) *Xbp1* mRNA (~ 2.2 fold) in rNLS8 mice at the 1-week timepoint. These findings further indicate the early activation of the ISR and broader UPR signaling from the beginning of induction of hTDP-43^ΔNLS^ expression in rNLS8 mice, with later loss of this potentially protective response during disease progression.

In line with RT^2^ array data, we confirmed the mRNA levels for five further upregulated genes that were significantly increased in the cortex of rNLS8 mice at 2-or 4-week timepoints compared to control mice (Fig 3C), including those involved in: neuroinflammation, being *CclI12*, *Il6*, *Tnf* and *Trl4*, and; oxidative stress response, being *Homx1.* Notably, the expression of these five genes was not significantly different between rNLS8 and control mice at the 1-week timepoint, suggesting that activation of neuroinflammation and the oxidative stress response are later events than ISR and apoptotic signaling in rNLS8 mice.

Concerning the two genes identified as downregulated in the array data, we verified the downregulation of *Ldha* and *Slc9a3* genes in the cortex of rNLS8 mice compared to the control mice at the 4-week timepoint, in line with the RT^2^ array data (Fig 3D). Unexpectedly, although both genes were not different between groups at the 2-week timepoint, we also detected a decrease of *Ldha* mRNA in the cortex of rNLS8 mice relative to control mice at the 1-week timepoint, suggesting early deficiency in metabolism may occur in rNLS8 mice. In contrast, we observed a significant *increase* of *Slc9a3* mRNA at the 1-week timepoint compared to control mice, suggesting that effects of disruption of sodium absorption may change over time in rNLS8 mice. Surprisingly, we detected a decrease of neuroinflammatory *Crp* mRNA in the cortex of rNLS8 mice at the 4-week timepoint in qPCR results, in contrast to the increase of *Crp* in the RT^2^ array, suggesting that changes in this gene may not be a consistent feature of disease in the rNLS8 mice (Fig 3D). Together, these findings reveal involvement of several important pathways, most notably activation of the ISR, beginning even prior to disease onset and persisting throughout early disease stages in rNLS8 mice.

To further biochemically validate activation of the ISR, we conducted immunoblotting (IB) to assess the phosphorylation of the eukaryotic initiation factor 2α (p-eIF2α) at serine 51 (S51) (Fig 3E), the core effector of the ISR, and the selective downstream target ATF4 (Fig 3H) (11), in rNLS8 mice at early disease stage (4 week timepoint). The IB results revealed slight but significant increases of eIF2α protein (~ 1.1 fold, Fig 3F) and significant increases of p-eIF2a (S51) (~ 1.4 fold, Fig 3G) in the cortex of rNLS8 mice compared to control mice at the 4 week timepoint. Notably, we also detected a dramatic increase of ATF4 protein (~ 3.8 fold, Fig 3I) in rNLS8 mice relative to control mice. These data further demonstrate activation of the ISR caused by accumulation of TDP-43 pathology in rNLS8 mice at early disease stages.

### Antisense oligonucleotide-mediated knockdown of Chop does not ameliorate motor deficits in rNLS8 mice

Apoptosis has been proposed as the final pathway leading to neurodegeneration in ALS/FTD (10), and CHOP is a key pro-apoptosis modulator involved in various cellular stress pathways relevant to disease, including ISR, oxidative stress and DNA damage (13). Our array and qPCR results revealed consistent upregulation of Atf4 and *Chop*, as well as *Bid*, which is an apoptotic gene downstream of Chop signaling, in the cortex of rNLS8 mice as early as the 1-week timepoint. We thus hypothesized that knockdown of *Chop*, as a possible early driver of apoptotic signaling in rNLS8 mice, would protect neurons against cell death and thereby prevent motor deficits. Previously, several RNase H-gapmer ASOs were reported to specifically target the *Chop* gene, successfully decreasing levels of *CHOP* protein in cells and mice (18).

We thus examined two of these previously reported *Chop* ASOs (18), and selected the most effective *Chop* ASO (previously reported as ASO#3) that suppressed *Chop* gene expression in the cortex of rNLS8 mice relative to control levels (Supplementary Fig. 2), for subsequent experiments in rNLS8 mice.

To investigate the effect of knockdown of *Chop* in rNLS8 mice, we conducted bilateral ICV injection of *Chop* ASO or its vehicle (saline) to adult littermate control or rNLS8 mice (10 weeks old), two weeks prior to Dox removal to ensure that decreased *Chop* mRNA occurred from the beginning of induction of *hTDP-43^ΔNLS^* expression, as shown in our preliminary studies (Supplemental Fig 2B, Fig 4A). We examined neurological and behavioural phenotypes, TDP-43 pathology and motor neurodegeneration in treated mice. Firstly, qPCR showed a significant increase (by ~1.5 fold) in *Chop* mRNA in the cortex of saline-treated rNLS8 mice (rNLS-Sal) at mid-disease stage of 6 weeks off Dox (eight weeks after ICV injection of ASOs), compared to the saline-treated control mice (Ctrl-Sal) (Fig 4B). The data indicates that the early increases in *Chop* mRNA previously detected from 1 week off Dox continued until at least this later disease state at 6 weeks off Dox. The qPCR data also confirmed that *Chop* ASO successfully decreased the *Chop* mRNA in the cortex in rNLS8 mice (rNLS-ASO), to approximately the level of control saline-injected mice (Fig 4B). Although there were no significant differences in the mRNA levels of *Chop* in the spinal cord between Ctrl-Sal and rNLS-Sal mice, *Chop* was decreased by ~40% in the spinal cord of the rNLS-ASO group compared to the Ctrl-Sal or rNLS-Sal groups (Fig 4B), indicating that direct ICV injection of *Chop* ASO can effectively decrease both cortex and spinal cord *Chop* gene expression.

**Fig. 4.**
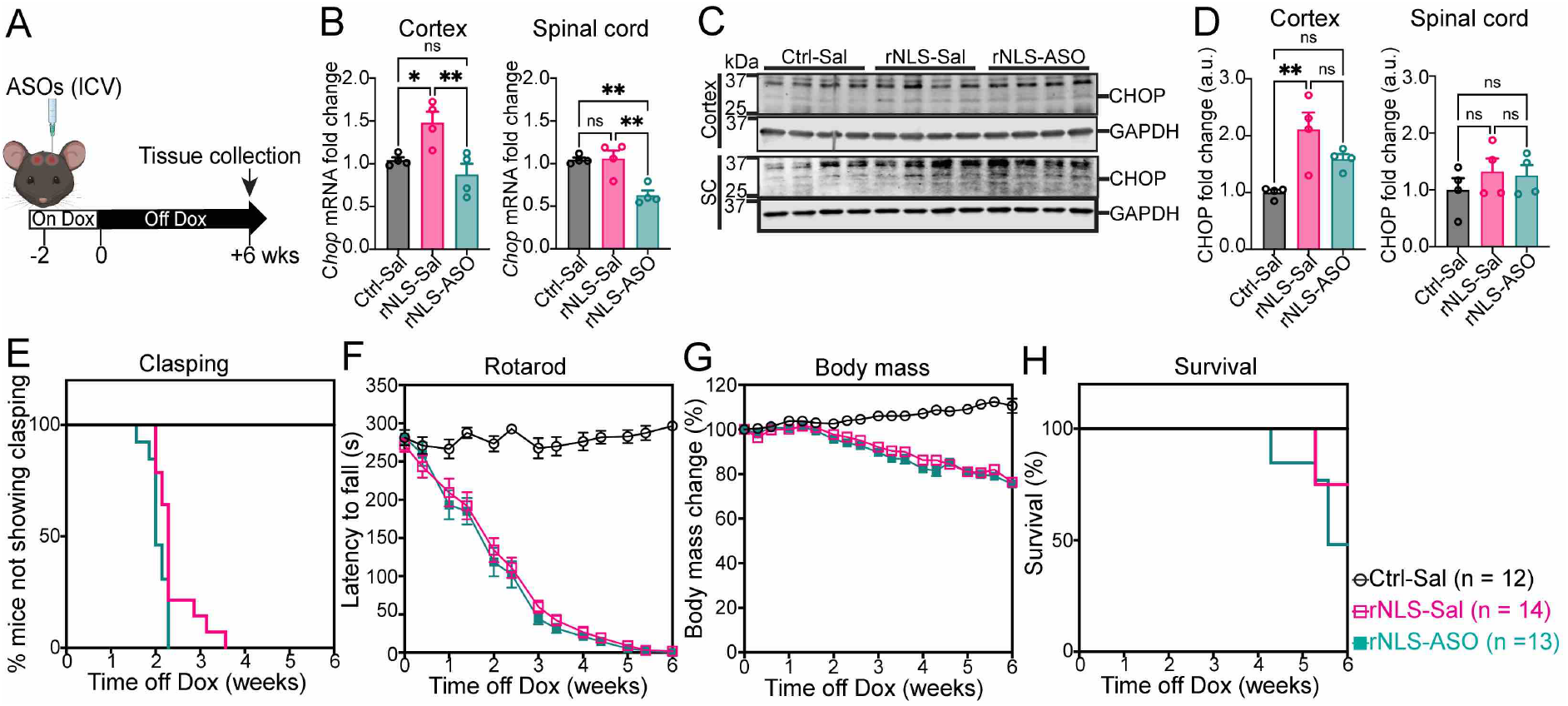
CHOP protein levels are increased in rNLS8 mice at 6 weeks off Dox, but ASO-mediated knockdown of Chop prior to induction of hTDP-43^ΔNLS^ expression does not alter disease phenotypes. **A**. Experimental schema. **B**. Real-time qPCR assessment demonstrated Chop ASO decreased Chop mRNA levels in the cortex and spinal cord of treated rNLS8 mice at 6 weeks off Dox (WOD) (n = 4). Data are normalised to Actb as the housekeeping gene. **C**. Immunoblotting for levels of CHOP protein in in the cortex and spinal cord of treated rNLS8 mice at 6 WOD (n = 4). Approximate molecular weights (kDa) are indicated. **D**. Quantification of immunoblots showed increased CHOP protein in the cortex and a trend of decrease of CHOP protein with ASO treatment in rNLS8 mice at 6 WOD. However, there were no significant differences between groups in CHOP protein levels in the lumbar spinal cord. **E-G**. Compared to saline-treated rNLS8 mice (rNLS-Sal), Chop ASO treatment led to no apparent beneficial effects on pathological phenotypes and motor deficits in clasping (**E**) or rotarod test (**F**), and did not alter the decline of body mass (**G**) or affect the survival of mice by the middle-disease stage at 6 WOD (**H**). n > 12. Mean + SEM. *p < 0.05 and **p < 0.01.

Furthermore, we conducted immunoblotting to analyse *CHOP protein* levels in the cortex and spinal cord of rNLS8 mice after ASO treatment (Fig 4C). Despite difficulties in detecting this protein via immunoblotting, we identified significant elevation of *CHOP protein* in the cortex of the rNLS8 mice (rNLS-Sal) of approximately 2.1-fold compared to the control (Ctrl-Sal), with a trend for a decrease in *CHOP protein* in rNLS-ASO mice (approximately 1.6-fold compared to rNLS-Sal mice), although this decrease did not reach statistical significance (Fig 4D). *CHOP protein* levels were difficult to detect in the spinal cord and there were no apparent differences in the levels of *CHOP protein* in the spinal cord of treatment groups despite the decrease detected via qPCR, although there was substantial variation in detected protein levels between mice (Fig 4D). Overall, consistent with previous evidence for a successful decrease of *CHOP protein* upon treatment with the ASO used here (18), our qPCR and IB results confirmed successful knockdown of *Chop* in the nervous systems of rNLS8 mice.

We further examined the effects of knockdown of *Chop* gene on neurological phenotypes of rNLS8 mice until late disease stages. rNLS8 mice developed characteristic features of progressive neurological decline compared to control mice, including hind-limb clasping onset (Fig 4E), progressive motor dysfunction (Fig 4F), body weight loss (Fig 4G), and lifespan (Fig 4H), similar to results previously reported (19). Notably, *Chop* ASO treatment had no significant effects on any of the examined neurological phenotypes (Fig 4E–4H). Further, rNLS-ASO mice also showed no difference in survival within 6 weeks off Dox compared to the rNLS-Sal mice (Fig 4H), although later disease stages were not assessed. Overall, these results demonstrate that knockdown of *Chop* did not affect the onset or early progression of motor phenotypes in rNLS8 mice.

### Knockdown of Chop does not alter overall TDP-43 pathology in rNLS8 mice

We next examined whether knockdown of *Chop* influenced TDP-43 pathology in the brains and spinal cords of rNLS8 mice. We detected significant increases of total TDP-43 protein (human and mouse TDP-43) in the RIPA-soluble fractions of cortex (Fig 5A, 5B) and spinal cord (Fig 5E, F) of rNLS-Sal mice compared to the Ctrl-Sal group, as expected. However, *Chop* knockdown caused no significant changes to the levels of RIPA-soluble total TDP-43 protein in cortex or spinal cord of rNLS8 mice compared to the rNLS-Sal group (Fig 5A, 5B, 5E, and 5F). The levels of total TDP-43 protein were also dramatically increased in the RIPA-insoluble fraction of both cortex (Fig 5A, 5C) and spinal cord (Fig 5E, 5G) of the rNLS-Sal mice relative to Ctrl-Sal mice, as seen previously (19). Notably, we detected accumulation of RIPA-soluble and-insoluble TDP-43 C-terminal fragments (CTFs) in the rNLS8 mice with the antibodies and conditions used in this study (Fig 5A, 5E). *Chop* knockdown had no effect on levels of total, CTF or high-molecular weight TDP-43 in the RIPA-insoluble fraction in the cortex of rNLS8 mice (Fig 5A, 5C), and there was similarly no effect on total or the high-molecular weight TDP-43 species in the RIPA-insoluble fraction in the spinal cord of rNLS8 mice (Fig 5E, 5G). However, interestingly, the spinal cord samples from rNLS-ASO mice showed a modest but statistically significant decrease in the low-molecular-weight TDP-43 CTFs (20-37 kDa) compared to rNLS-Sal mice (Fig 5G). We further examined levels of pTDP-43 at serine (409/410) in the RIPA-insoluble protein fractions, demonstrating significant increases of pTDP-43 in both cortex (Fig 5D) and spinal cord (Fig 5H) of the rNLS-Sal mice compared the Ctrl-Sal mice, as expected. Nevertheless, knockdown of *Chop* did not change the levels of pTDP-43 in either cortex or spinal cord of rNLS8 mice.

**Fig. 5.**
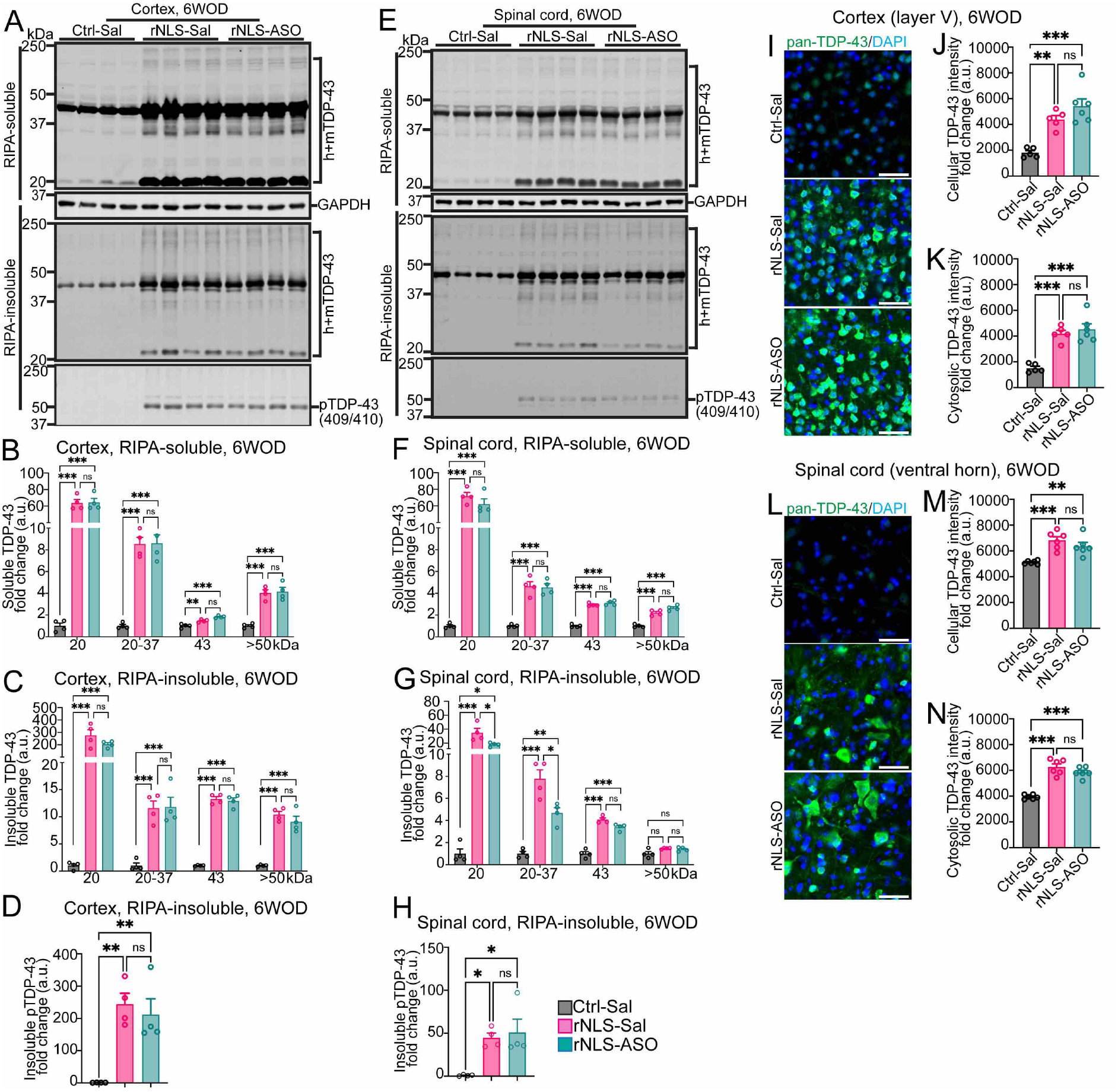
ASO-mediated knockdown of Chop has little effect on TDP-43 levels or solubility in the cortex and spinal cord of rNLS8 mice at 6 weeks off Dox. IB assessed TDP-43 in the RIPA-soluble and RIPA-insoluble fractions and p-TDP-43 in the RIPA-insoluble fractions for the cortex (**A-D**) and spinal cord (**E-H**) of mice at 6 weeks off Dox (WOD) (n = 4). Approximate molecular weights (kDa) are indicated on the left and target protein on the right. GAPDH was used as the loading control for RIPA-soluble IB and total protein for RIPA-insoluble IB. Quantification of IB for different molecular weight species of TDP-43 or p-TDP-43 in the RIPA-soluble and-insoluble fractions in the cortex (**B-D**) and the spinal cord (**F-H**) (n = 4). Representative IF images for pan-TDP-43 staining (human + mouse) (green) with the nuclear marker DAPI (blue) in the cortex (**I**) and spinal cord (**L**) of treated mice at 6WOD. Scale bars as 50 μm. Quantification of IF for pan-TDP-43 protein in identified cell or cytoplasmic region in the cortex (**J, K**) and spinal cord (**M, N**) of treatment groups. Ctrl-Sal (n = 5), rNLS-Sal (n = 6) and rNLS-ASO (n = 6). Mean + SEM. *p < 0.05, **p < 0.01 and ***p < 0.001.

To further analyse the effect of *Chop* knockdown on TDP-43 accumulation in the cytoplasm of neurons in rNLS8 mice, we conducted IF for TDP-43 and analysed neurons in the cortex (Fig 5I) and spinal cord (Fig 5J), with TDP-43 primarily localised in the nuclei of neurons of the Ctrl-Sal mice. In contrast, TDP-43 was localised to cytoplasmic regions in the neurons in the layer V of the cortex (Fig 5I), and neurons in the ventral horn of the lumbar spinal cord in rNLS8 mice (Fig 5L), as expected (19). Nevertheless, *Chop* ASO treatment did not result in any changes in overall levels or cytoplasmic TDP-43 distribution in neurons of cortex or spinal cord in rNLS8 mice. Quantification of IF confirmed the elevation of TDP-43 in the rNLS-Sal and rNLS-ASO mice shown as fluorescence intensity measured per cell or within the cytosolic regions in cortical (Fig 5J, 5K) and spinal cord neurons (Fig 5M, 5N). Nevertheless, suppression of *Chop* by ASO led to no significant differences in total or cytosolic fluorescence intensity of TDP-43 between rNLS-Sal and rNLS-ASO groups. Overall, these data demonstrate that knockdown of *Chop* did not consistently affect the overall accumulation or mislocalisation of TDP-43 in rNLS8 mice.

### rNLS8 mice display changes in apoptosis signaling over time and increased cleaved caspase-3 levels that are not affected by knockdown of Chop

CHOP protein acts as a critical transcription factor that directly regulates the expression of apoptosis genes, coordinating the upstream initiation of apoptosis proteins (13). However, it remains unclear how TDP-43 pathology affects the regulation of the downstream apoptotic mediators of CHOP activation. Thus, we first examined the expression of these genes in rNLS8 mice at early disease stages. The qPCR results revealed significant elevation of anti-apoptotic *Bcl2* gene (~ 1.3 fold, Fig 6A), Bcl2 homology 3 (BH3)-only pro-apoptotic initiator *Bim* gene (~ 1.5 fold, Fig 6B) and *Noxa* gene (~ 2.2 fold, Fig 6C) in the cortex of rNLS8 mice before disease phenotypes at 1 week off Dox, compared to control mice. There were no significant changes in *Bcl2* or *Bim* at 2 or 4 weeks off Dox in rNLS8 mice, although upregulation of *Noxa* was detected at 2 weeks but not 4 weeks in rNLS8 mice (Fig 6C). Notably, *Puma* was not altered at 1 week off Dox but was significantly increased in the rNLS8 mice at 2-and 4-week timepoints, after disease onset. Dysregulation of these apoptosis genes suggests involvement of apoptosis signaling in neurodegeneration caused by TDP-43 pathology in rNLS8 mice, and suggest that there is differential switch from early anti-apoptotic to later pro-apoptotic signaling over the disease course.

**Fig. 6.**
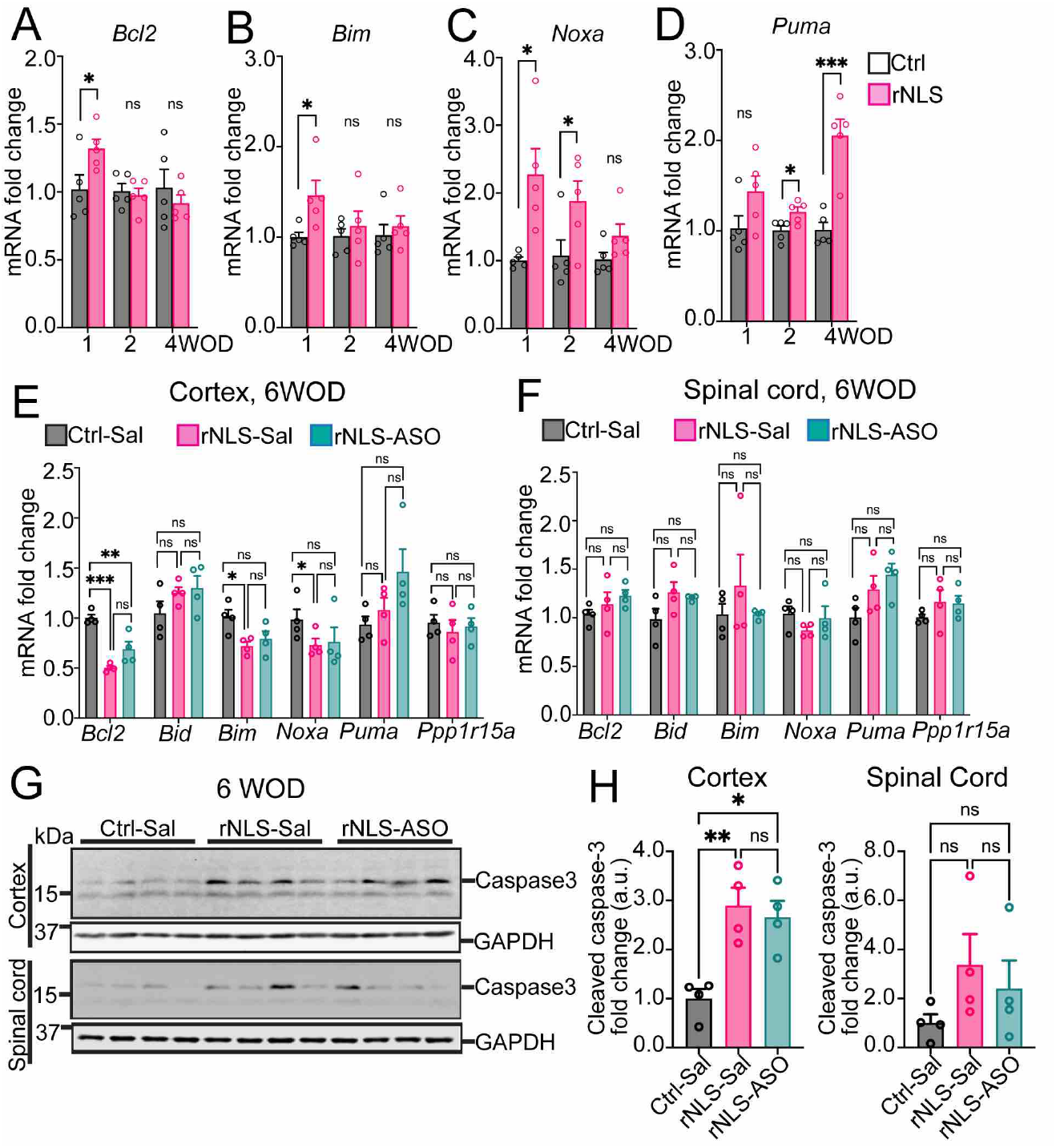
rNLS8 mice display early activation of apoptosis, and increased cleaved caspase-3 levels in the cortex at 6 weeks off Dox that is not altered by knockdown of Chop. Real-time qPCR results revealed dysregulation of Chop target apoptosis genes in the cortex of rNLS8 mice at 1, 2 and 4 weeks off Dox (WOD) (n = 5): **A**. Bcl2; **B**. Bim; **C**. Noxa; **D**. Puma. Real-time qPCR results revealed expression changes of Chop downstream genes in (**E**) and spinal cord (**F**) of mice at 6 WOD after ASO treatment (n = 4). Actb as the housekeeping gene. **G**. IB for caspase-3 in the cortex and spinal cord of mice at 6 WOD (n = 4). Approximate molecular weight markers (kDa) are shown, and GAPDH is used as loading control. **H**. Quantification of IB for cleaved caspase-3 (17 kDa) in the cortex and spinal cord of treated mice. Mean + SEM. *p <0.05, **p < 0.01 and ***p < 0.001.

Further, we assessed the expression of these apoptotic genes in rNLS8 mice at the mid-stage of disease (6 weeks off Dox) and with knockdown of Chop. Interestingly, we detected significantly decreased mRNA levels of the anti-apoptotic *Bcl2* gene as well as pro-apoptotic *Bim* and *Noxa* genes, and no difference in *Puma* levels, in the cortex of the rNLS-Sal mice compared to the Ctrl-Sal mice at 6 weeks off Dox (Fig 6E). This is in contrast to the upregulation of these genes identified at earlier disease stages of rNLS8 mice, and may be due to extensive neurodegeneration that occurs at this stage of disease (19). Notably, *Bid* expression, which was upregulated in rNLS8 mice at early disease stages (1,2 and 4 weeks off Dox, Fig 3A), was not significantly different between control mice (Ctrl-Sal) and rNLS8 mice at this later disease stage (6 weeks off Dox). rNLS8 mice also displayed no alteration of examined genes in the spinal cord relative to the control mice (Ctrl-Sal) at 6 weeks off Dox (Fig 6F). Further, *Chop* knockdown had no effect on the mRNA levels of anti-or pro-apoptosis genes in either cortex or spinal cord of treated mice at 6 weeks off Dox (Fig 6E, 6F).

Given that caspases play essential roles in orchestrating apoptosis and caspase-3 is a primary downstream executioner among these proteases, we also examined the activation of caspase-3 in the cortex of rNLS8 mice by quantifying the levels of a specific 17 kDa caspase-3 cleavage product (Fig 6G), which is required for caspase-3 activity in apoptosis (21). We detected a significant elevation of cleaved caspase-3 (~17 kDa band) in the cortex of the rNLS8 mice (rNLS-Sal) compared to the control mice (Ctrl-Sal) at the 6-week timepoint (Fig 6H), suggesting that activation of apoptosis may drive neurodegeneration in the later stages of disease in rNLS8 mice. Similarly, there was a trend of increased cleaved caspase-3 levels in the spinal cord of rNLS8 mice (rNLS-Sal) relative to the control mice (Ctrl-Sal), although it did not reach statistical significance (Fig 6H). Elevation of caspase-3 is consistent with the previous report that demonstrated dramatic motor neuron loss in the cortex of rNLS8 mice at 6 weeks off Dox (19). Nevertheless, knockdown of *Chop* had no effect on the level of cleaved caspase-3 in either cortex or spinal cord of treated mice (rNLS-ASO) relative to the rNLS-Sal group (Fig 6H). Overall, these data demonstrate that knockdown of *Chop* did not ameliorate the disease-associated activation of apoptosis signaling in rNLS8 mice, suggesting that alternative pathways or modulators may account for the regulation of these genes during apoptosis caused TDP-43 pathology.

### rNLS8 mice display dramatic astrogliosis and dysregulation of astrocytic genes, which are not ameliorated by knockdown of Chop

In addition to the cell-autonomous effect of ISR in neurons, dysfunctional astrocytes can also contribute to the pathogenesis of human neurodegenerative diseases (22). Given that astrogliosis is an established feature of both ALS (23) and the rNLS8 mouse model (19), we assessed the activation of astrocytes that contribute to glutamate-induced motor neuron death in ALS (24) and which would be expected to be ameliorated by a beneficial treatment strategy. Firstly, we assessed astrocytes using the pan-reactive astrocytic marker GFAP by IF (Fig 7A, 7D). The results indicated a dramatic increase in the number of GFAP-positive astrocytes with ramified cell processes in the cortex and spinal cord of rNLS8 mice (rNLS-Sal) compared to the Ctrl-Sal mice (Fig 7B). Knockdown of *Chop* slightly but significantly increased the number of reactive astrocytes in treated mice (rNLS-ASO) relative to that of rNLS-Sal mice (Fig 7B), with a slight elevation of GFAP intensity in the cortex of rNLS-ASO mice compared to the control mice (Ctrl-Sal) (Fig 7C). However, we found no difference in either astrocyte number (Fig 7E) or GFAP intensity (Fig 7F) in the spinal cord among treatment groups.

**Fig. 7.**
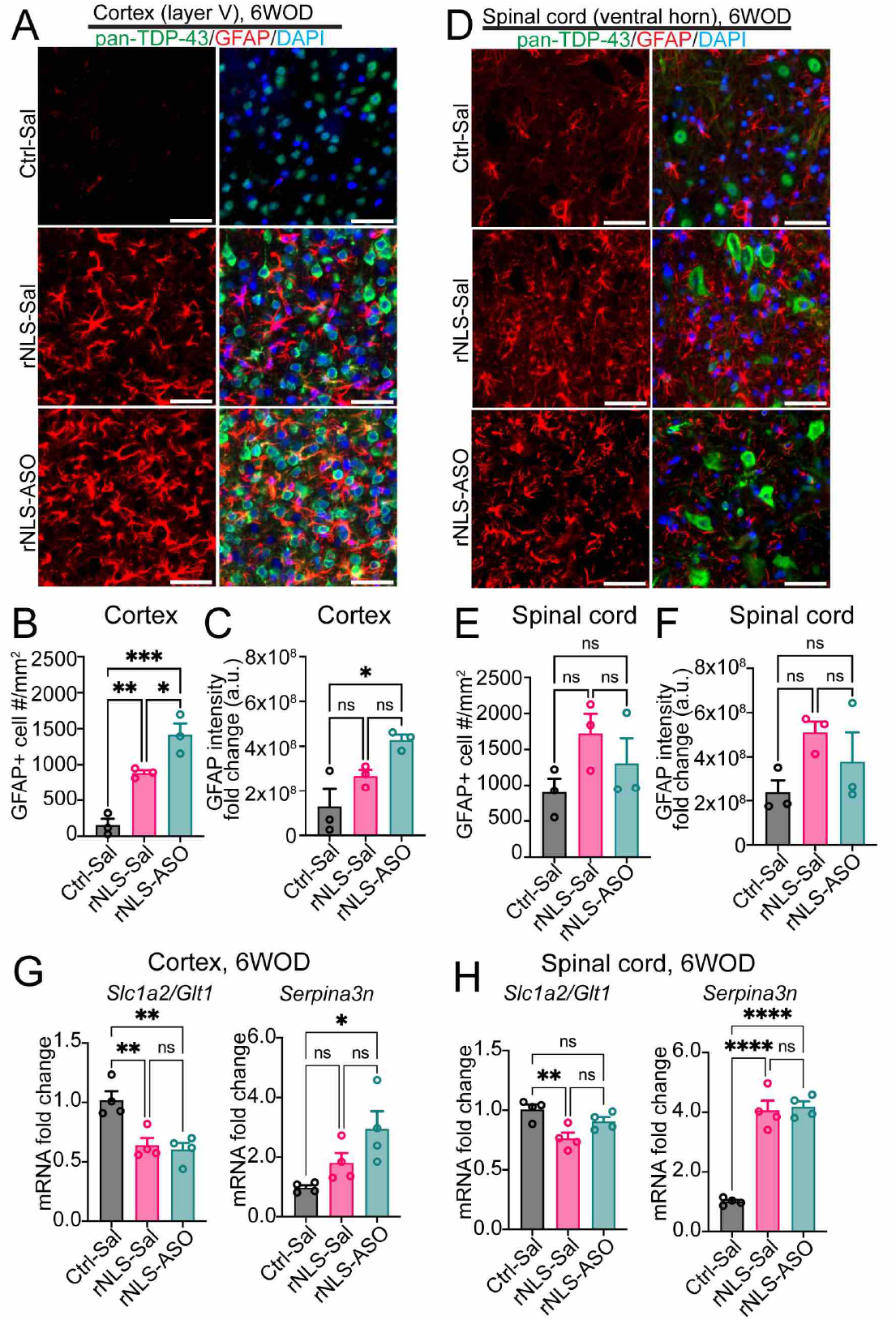
rNLS8 mice develop reactive astrogliosis and altered astrocytic gene expression, which are not ameliorated by knockdown of Chop. Representative IF images for pan-TDP-43 (human + mouse) (green), reactive-astrocytic marker GFAP (red), and the nuclear marker DAPI (blue) in the layer V of cortex (**A**) or the ventral horn of spinal cord (**D**) of mice at 6 weeks off Dox (WOD). Scale bars represent 50 μm. Cell counting for GFAP positively labeled astrocytes in the cortex (**B**) and spinal cord (**E**) of treated mice (n = 3). Quantification of IF for GFAP intensity in the cortex (**C**) and spinal cord (**F**) of treated mice at 6 WOD (n = 3). qPCR analysis of astrocytic gene expression in the cortex (**G**) and spinal cord (**H**) of Chop ASO-treated mice at 6 WOD (n = 4). Data are normalised to Actb as the housekeeping gene. Mean + SEM. *p < 0.05, **p < 0.01, ***p<0.001 and ****p < 0.0001.

The ISR has been shown to impair the neurotrophic function and to modulate inflammatory states of astrocytes, by dysregulation of astrocytic gene expression (25). Therefore, to determine whether knockdown of *Chop* affected the expression of genes important for astrocytic physiological function and reactivity, we examined levels of two representative astrocytic genes. The astrocytic solute carrier family 1 member 2 gene *Slc1a2/Glt1* that encodes glutamate transporter (GLT1/EAAT2) plays key roles in glutamate reuptake and modulation of homeostatic brain function, and is deficient in ALS (26). The pan-reactive astrocytic gene *Serpina3n* encodes the serine protease inhibitor that is upregulated during neurodegeneration (22). We detected a significant decrease of the *Slc1a2/Glt1* mRNA in the cortex (Fig 7G) and spinal cord (Fig 7H) of rNLS8 mice (rNLS-Sal) compared to the control mice (Ctrl-Sal). However, there were no significant differences in the *Slc1a2/Glt1* levels of rNLS-ASO mice relative to the rNLS-Sal group in the cortex or spinal cord. Similarly, rNLS8 mice (rNLS-Sal) displayed dramatic upregulation of *Serpina3n* in the spinal cord compared to Ctrl-Sal mice (Fig 7H), although *Serpina3n* was not significantly different in the cortex between Ctrl-Sal and rNLS-Sal mice (Fig 7G). Nevertheless, knockdown of *Chop* did not affect *Serpina3n* levels in treated mice (rNLS-ASO) compared to the rNLS-Sal group. Overall, these results not only confirmed previous reported astrogliosis in rNLS8 mice at the mid-disease stage (19) but also revealed dramatic dysregulation of disease-related astrocytic genes, suggesting disruption of astrocytic function may contribute to neurodegeneration in this model. Nevertheless, we did not identify any evidence for amelioration of disease-related astrogliosis or astrocytic gene expression alterations upon knockdown of *Chop* in rNLS8 mice.

## Discussion

Here, we aimed to determine which cellular stress pathways contribute to disease onset and progression in ALS, using a well-characterised mouse model of cytoplasmic TDP-43 proteinopathy. Our data identified early dysregulation of multiple genes in several signaling pathways in the cortex of pre-onset rNLS8 mice, including *Atf4*, *Chop*, *Bid*, *Gadd45g* and *Trp53*, extending throughout disease progression. Most notably, we identified early activation of the ISR and alterations in apoptosis pathways even in rNLS8 mice prior to onset of motor phenotypes and continuing through early disease stages, leading to activation of caspase-3. However, suppression of *Chop*, a key activator of the ISR, beginning prior to induction of hTDP-43^ΔNLS^ expression, did not ameliorate disease pathology, onset or progression in rNLS8 mice. Together, these studies demonstrate that cellular stress pathways, possibly including early anti-apoptotic signaling, are activated early prior to disease onset and may contribute to neurodegeneration in rNLS8 mice. However, suppression of *Chop* throughout disease is not sufficient to prevent a switch to pro-apoptotic signaling associated with disease progression.

In contrast to healthy neurons with tightly controlled protein synthesis, folding and degradation, in disease protein aggregation triggers the ISR which can ultimately lead to neuronal death (11, 12). TDP-43 pathology is associated with activation of ISR in both neurons and non-neuron cells in nervous system postmortem tissues of people who have died with ALS or FTD (27), with increased *Chop* levels identified in human ALS (17) as well as in mouse models for ALS/FTD (15, 18, 28, 29). *In vivo*, TDP-43 pathology also results in global protein translation inhibition, which is the main effect of the ISR (30). Moreover, in cell culture, disease-associated mutant TDP-43 proteins trigger ISR activation and elevation of downstream gene expression including *Chop,* involving an amplifying cell stress response potentially leading to cell death (15). *Chop/Ddit3* was also recently identified as one of the top upregulated genes in a human neuronal model of TDP-43 pathology, and was shown to be a direct target of TDP-43-mediated gene expression with increased *Chop* levels in neurons with depleted nuclear TDP-43 in human FTLD-TDP-derived samples (31). In addition, mice expressing ALS-associated mutant SOD1 (29, 32) or ALS/FTD-associated FUS (33) also display activation of ISR and/or impairment of ongoing protein synthesis, including upregulation of *Chop* (34). The ISR is also activated in neurons expressing expanded C9orf72 hexanucleotide repeats, and is similarly seen in the cortices of people who have died with C9orf72-linked ALS/FTD (35). Together, these studies indicate that the ISR, and *Chop* in particular, is involved broadly in disease, although it remains unclear whether these pathways may be amenable to therapeutic targeting in TDP-43-related disease. Our data show that knockdown of *Chop* using ASO decreased *Chop* expression (Supplementary Fig 2), but had no impact on overall TDP-43 pathology (Fig 5) or motor deficits (Fig 4). These findings are consistent with previous reports that neither genetic knockout nor ASO-mediated knockdown of *Chop* led to beneficial outcomes in the mutant SOD1^G93A^ mouse model for ALS (18, 36) or rTg4510 mice for Alzheimer’s disease (37).

Prolonged ISR activation induces apoptosis via cell death downstream of the intrinsic pathway that is regulated by Bcl2 family members (*Bim*, *Noxa*, *Puma*), and the extrinsic pathway (*Bid*) that can be triggered by various cell death signals (10). Notably, our results revealed that induction of hTDP-43^ΔNLS^ expression rapidly induced both anti-apoptotic (*Bcl2*) and pro-apoptotic genes (*Bim, Bid, Noxa*) in rNLS8 mice even before disease onset (Fig 6). In contrast, from disease onset, *Bcl2* mRNA in rNLS8 mice returned to the level similar to the control mice (Fig 6E), coinciding with elevation of pro-apoptotic *Bid*, *Noxa* and *Puma* mRNAs in rNLS8 mice, suggesting a switch from pro-survival to pro-death signalling over time. Importantly, our findings revealed that activation of these pro-apoptotic genes occurred differentially over the disease course (Fig 6). For example, *Bim* mRNA was increased only prior to disease onset, in line with the previous report that *Bim* gene can be induced by overexpression of TDP-43 in neurons (28). In contrast, p53-mediated transcription of *Noxa* and *Puma* was increased in rNLS8 mice during early disease stages, consistent with our data showing upregulation of *Trp53* and *Gadd45g* (Fig 3A). Notably, *Bid*, a gene that can be induced by a variety of upstream signals from the ISR, DNA damage, oxidative stress and death receptors (for example, TNF and its receptor), was upregulated prior to disease onset and into early disease stages. This differential regulation of apoptotic genes indicates that multiple stress signaling pathways may co-contribute to and even enhance pro-apoptotic signaling in rNLS8 mice at different disease stages. Moreover, we speculate that significant neuron loss occurs in the motor cortex of rNLS8 mice by 6 weeks off Dox, which may account for the late decrease of *Bcl2*, *Bim* and *Noxa* expression in rNLS8 mice (Fig 6E), consistent with previous findings that show decreases of *Bcl2* mRNA in postmortem tissues in ALS and other human neurodegenerative diseases (21). Importantly, rNLS8 mice displayed elevation of pro-apoptotic executioner caspase-3 (Fig 6H) at the mid-stage of disease, providing direct evidence that supports the previous findings of increased caspase-3 in brain and spinal cord autopsy tissues of people with ALS (38). Overall, our data therefore supports the hypothesis that TDP-43 pathology induces apoptosis contributing to neurodegeneration, but also reveals complex regulation between pro-survival and prodeath signaling that begins early in the disease course. Thus, targeting apoptosis may be a beneficial therapeutic strategy for treatment of ALS/FTD. Notably, we did not detect any change in expression of necroptosis genes *Ripk1* and *Ripk3* at 2 or 4 weeks off Dox in rNLS8 mice in the RT^2^ array, despite the detected alterations of apoptosis signalling even at these early timepoints, aligning with recent work suggesting that necroptosis is not a primary driver of neurodegeneration in ALS (39).

Our results also demonstrated dramatic astrogliosis in the cortex and spinal cord of rNLS8 mice (Fig 7), with significant upregulation of pan-reactive astrocytic marker *Serpina3n* in rNLS-ASO mice, which can be induced upon neuronal damage and contributes to neuroinflammation [30, 32]. Our finding is consistent with previous studies showing upregulation of *Serpina3n* in a conditional TDP-43 knockout mouse model for FTD (40), suggesting that the loss of endogenous nuclear TDP-43 in neurons of the rNLS8 mice (19) may account for this phenotype. Notably, our results also indicated the downregulation of the *Slc1a2* gene encoding glutamate transporter GLT1 in rNLS8 mice (rNLS-Sal) (Fig 7G, 7H). This is in line with the previous observation that GLT1 expression is selectively decreased in astrocytes of ALS postmortem tissues (26) and in TDP-43 associated FTD cases (41). Hence, our results further support the hypothesis that disease-associated astrocyte reactivity and deficiency of astrocytic reuptake of glutamate at the synaptic cleft contribute to neurotoxicity in ALS.

In addition to ISR, other biological processes also appear active early in the rNLS8 mice, such as DNA damage response, unfolded stress response (UPR) and dysfunction in cellular metabolic functions. Our data revealed dramatic elevation of *Trp53* and *Gadd45g* genes as early as 1 week off Dox and continued to early disease stages (Fig 3A). Previous research show depletion of TDP-43 or ALS-linked mutant TDP-43 induces DNA damage and consequently p53-dependent apoptosis of motor neurons (42, 43), while GADD45G acts as a stress sensor for p53-mediated apoptosis (44, 45). Moreover, it is proposed that oxidative stress is a pathogenic mechanism in disease, in parallel with or downstream to ISR [4, 35]. Our results reveal elevation of the oxidative stress-responsive gene *Homx1* in the cortex of rNLS8 mice from the 2 week time point (Fig 3C), in line with the evidence from the motor cortex of ALS-linked mutant TDP-43 transgenic mice (6, 46, 47) and in the lumbar spinal cord of SOD1^G93A^ mice for ALS (48). However, most oxidative stress genes remained unchanged in our study, suggesting that further studies are required to identify the roles of oxidative stress genes in TDP-43-related disease. Additionally, we detected significant increases of the spliced (active) form of *Xbp1,* a key component of the UPR mediated by activation of IRE1, in rNLS8 mice at 1 and 2 weeks off Dox (Fig 3B). These results indicate involvement of early activation of UPR signaling in rNLS8 mice in response to accumulation of TDP-43 pathology, in addition to the ISR. Further, our data demonstrate early deficiency of cellular metabolism in rNLS8 mice at early disease stage (4 weeks off Dox). LDHA, encoded by the *Ldha* gene, is specifically expressed in cortical neurons (including Layer V neurons) in human and rodent brains, and plays a vital role in neuronal health, with knockdown of *Ldha* inducing intrinsic apoptotic (49–51). This suggests another mechanism by which apoptosis may be triggered in rNLS8 mice. More importantly, as many of these pathways are also regulated at translational and post-translational levels, it is worth further investigation using high-throughput protein screening, for instance, quantitative proteomics (52), to characterise the interplay among pathways involved in ALS and FTD.

Of relevance for therapeutic development, ASOs have become an increasingly promising class of drugs to target conventionally undruggable molecules or pathways in a broad range of human neurological and psychiatric diseases. For instance, the FDA-approved Nusinersen targeting Smn2 to treat spinal muscular atrophy has shown remarkable efficacy in the clinic (53). Recently, there have been newly developed ASO drugs for ALS/FTD in clinical trials, such as Tofersen to decrease SOD1 protein synthesis (54) and an ASO to inhibit translation of C9orf72 mRNA (WVE-004, Wave Life Sciences). Moreover, preclinical trials have also been undertaken to ameliorate TDP-43 pathology via modulation of TDP-43-regulating proteins such as *Ataxin-2* [63, 64], which are now in Phase I clinical trials (ClinicalTrials.gov, Identifier# NCT04494256). Further identification of disease-relevant targets that are able to protect against neurodegeneration caused by TDP-43 dysfunction offers hope for development of additional improved therapeutics, potentially including antisense approaches, for ALS and FTD (55).

### Limitations

Among the limitations of our work, we analysed a single timepoint of administration of the *Chop* ASO, namely two weeks prior to the removal of Dox from rNLS8 mice. This allowed us to assess the effects of *Chop* suppression from before induction of hTDP-43ΔNLs expression and throughout the entirety of the study. Nevertheless, early activation of ISR may be protective for cells under acute cellular stress conditions (11, 12, 16), and so it is possible that the early activation of *Chop* in rNLS8 mice is a beneficial response, at least early in the disease course. Indeed, our findings of shifts in activation or inhibition of different signalling pathways prior to onset and throughout disease indicate that the effects of ISR activation may be modulated by other contributing factors, such as changes in apoptosis signalling, at different disease points. Knockdown of *Chop* from the beginning of induction of hTDP-43^ΔNLS^ may have thereby ablated any neuroprotective effects of an initial ISR activation in neurons. Therefore, future studies could determine whether *Chop* suppression beginning only later in the disease course is beneficial in extending survival. Indeed, in this study we did not examine whether knockdown of *Chop* can modify the disease phenotypes or lifespan of rNLS8 mice beyond 6 weeks off Dox, which may be necessary to identify any beneficial longer-term effects. Furthermore, recent work has highlighted the importance of the level of cellular stress for effective responses to small molecule modulators of the ISR. For example, the small molecular ISRIB, which overcomes eIF2a phosphorylation-mediated inhibition of translation by stimulating eIF2B activity, effectively rescues protein translation inhibition under conditions of acute viral infection with low levels of p-eIF2a but is ineffective in chronic infection with high levels of p-eIF2a present (56). These results suggest that a critical threshold of ISR activation exists in disease conditions above which normally protective therapeutic approaches become ineffective (56). Additional studies are also required to investigate changes at a single-cell level. Regardless, our work also highlights other likely mediators of neurotoxicity which may be more promising targets for further development, including a switch between the balance or anti-and pro-apoptotic signalling. Further work is required to investigate therapeutically relevant targets and to define the timing of therapeutic intervention to successfully ameliorate disease.

### Conclusion

Taken altogether, we identified early activation of several distinct cellular stress pathways, notably the ISR and pro-apoptotic signalling, in the cortex of rNLS8 mice even prior to disease onset. These pathways are essential for neuronal survival-and-eath decisions, indicating that the biological pathway towards neurodegeneration begins very early after accumulation of cytoplasmic TDP-43 in disease. For the first time, our data revealed that cytoplasmic TDP-43 accumulation induces pro-apoptotic caspase-3 in parallel with decreases of anti-apoptotic Bcl2 in the central nervous system of TDP-43 mice (Fig 8). We conclude that multiple cellular stress pathways are active early in rNLS8 mice, notably the pro-apoptotic signalling that is activated even prior to disease onset in rNLS8 mice, which likely triggers eventual motor neuron death via apoptosis following loss of competing anti-apoptotic signals. Although knockdown of the key ISR mediator *Chop* had no effect on motor deficits and did not ameliorate overall TDP-43 pathology in rNLS8 mice, our findings indicate that targeting of cell stress and death signalling may be a potential promising avenue for treatment of TDP-43-associated ALS and FTD.

**Fig. 8.**
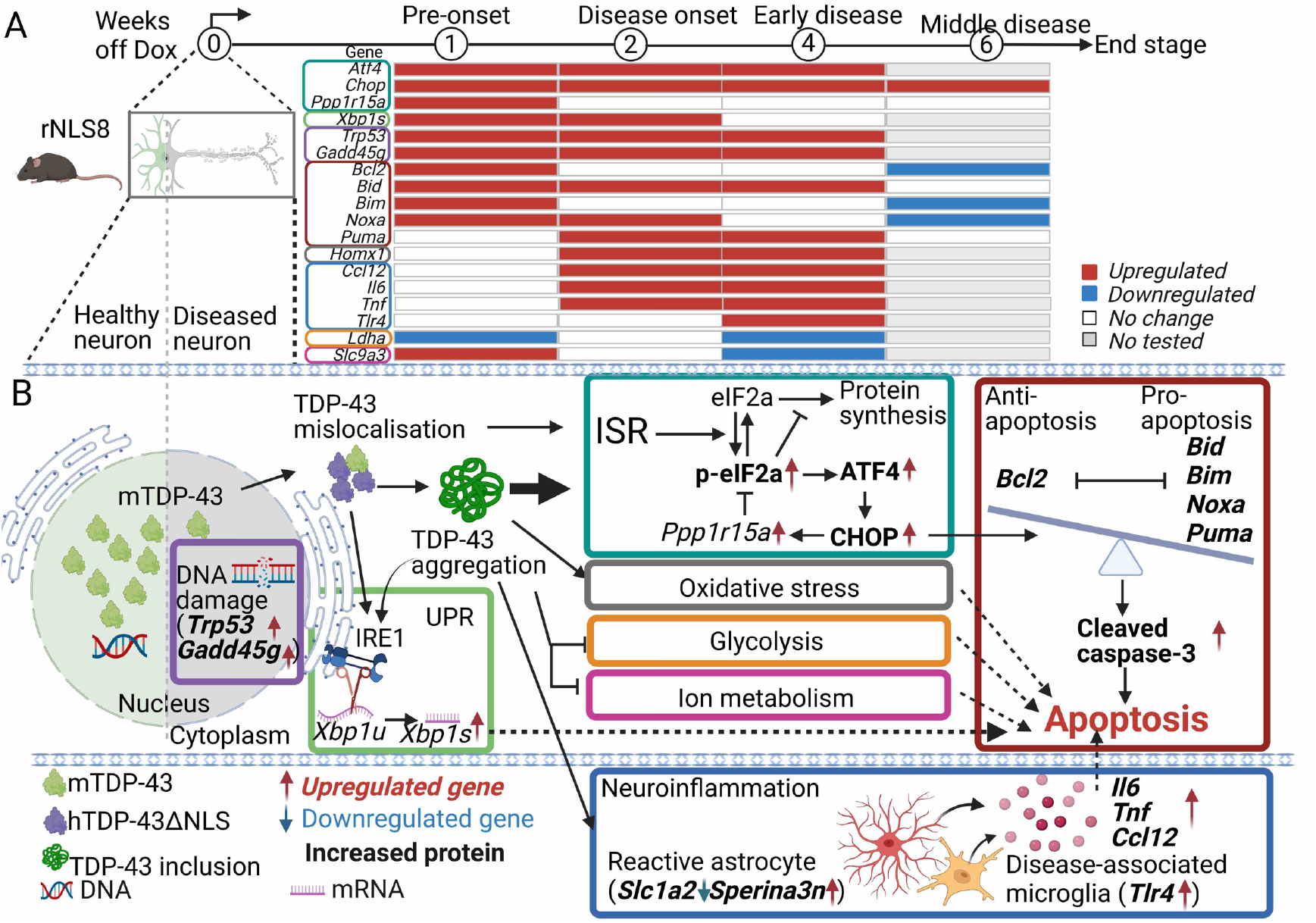
TDP-43 pathology causes early activation of the ISR and apoptosis signaling in rNLS8 mice. **A**. Summary table for gene expression changes in the cortex of rNLS8 mice over time. **B**. Mechanism schema. TDP-43 mislocalisation induces early activation of multiple cell stress signaling pathways in the cortex rNLS8 mice even before disease onset, such as the ISR, UPR, DNA damage response and apoptosis. Notably, the prolonged activation of these cell stress pathways caused by TDP-43 pathology, in particular the ISR, continues to early-disease stages where rNLS8 mice displayed dramatic elevation of p-eIF2α and ATF4 protein, and impairment of cellular metabolism (glycolysis and ion metabolism). Dysregulation of stress pathways lead to reduction of anti-apoptotic Bcl2 mRNA, continued increase of pro-apoptotic gene expression, and increases of cleaved caspase-3 and astrogliosis, likely contributing to neurodegeneration in rNLS8 mice.

## Materials and Methods

### Animals

To generate samples for RT^2^ qPCR arrays, rNLS8 transgenic mice and littermate controls on a mixed B6/C3H F1 background were produced as previously described (57). Founder monogenic B6;C3-Tg(NEFHtTA)8Vle/J (NEFH-tTA line 8, stock #025397) mice and monogenic B6;C3-Tg(tetO-TARDBP*)4Vle/J (tetO-hTDP-43ΔNLS line 4, stock #014650) mice were obtained from the Jackson Laboratory (Bar Harbor, ME, USA) (19). Intercross breeder and experimental mice were fed with chow containing 200mg/kg doxycycline (Dox) (Specialty Feeds, Australia). Experiments were conducted with approval from the Animal Ethics Committee of Macquarie University (#2016-026). For all other experiments, rNLS8 mice were produced from the intercross of homozygous tetO-hTDP-43ΔNLS line 4 mice with hemizygous NEFH-tTA line 8 mice both on a pure C57BL/6JAusb background following >10 generations of backcrossing and were fed with Dox-containing chow (200mg/kg, Specialty Feeds, Australia) (58). Experiments were conducted with approval from the Animal Ethics Committee of The University of Queensland (#QBI/131/18). During all experiments, mice were switched to normal chow to induce expression of hTDP-43ΔNLS. Male and female mice at approximately ten weeks of age were housed in temperature-and humidity-controlled conditions (21 ± 1 °C, 55 ± 5%) with a 12-h light/dark cycle (lights on at 06:00 h). Mice were randomly allocated to groups for all experiments of time point analyses and ASO administration groups, the size of which were calculated by G*Power (version 3.1). Both sexes were included and balanced between groups. Littermate non-transgenic and monogenic animals were used as controls. One mouse was excluded from all analyses due to non-neurological disease (malocclusion). Three mice in the ASO experiments were excluded from the qPCR and histological analyses since they reached the humane end-point prior to the predefined time point for tissue collection of 6 weeks off Dox. All experimental procedures were conducted under the guidelines of the National Health and Medical Research Council of Australia in accordance with the Australian Code of Practice for the Care and Use of Animals for Scientific Purposes.

### Surgery procedures and ASO administration

Two *Chop* ASOs were synthesised and purified as previously described (as per (#3 and #5) (18). For intracerebroventricular (ICV) injection, mice were anaesthetised by isoflurane (Abbott Laboratories) using an isoflurane machine (Kent Scientific Co. Torrington, CT, USAs) at the rate of 2-4% according to body weights of mice. Bilateral ICV injections were conducted stereotaxically using a 30 G needle connected to a Hamilton syringe by polyethylene tubing into lateral ventricles: AP = 0 (Bregma); L = + 0.9; depth = 2.3 (coordinates are in millimetres relative to the Bregma) (59). A total of 10 μL of ASO solution (50 mg/mL prepared in sterile saline solution) or saline solution as vehicle was injected into each of the left and right ventricles over 2 min (5 μL in each ventricle). After injections, mice were housed in home cages for two weeks before the removal of the Dox diet to induce neuronal hTDP-43^ΔNLS^ expression.

### Mouse monitoring and behavioural tests

Mice were monitored and weighed three times per week after Dox feed was removed as previously described (19, 57). Briefly, for observation of collapsing splay or clasping of hindlimbs, mice were suspended by the tail for >5 seconds. The failure to extend both hindlimbs was recorded as a positive response of collapsing splay and holding hindlimbs together was recorded as a presence of clasping splay. Mice were tested for rotarod performance of motor coordination and balance once per week, as previously described (19). Briefly, mice were placed on a rotarod apparatus (Ugo Basile SRL, Gemonio, Italy) at a speed of 5 rpm with acceleration up to 40 rpm within 300 s. The time to fall was recorded. If mice were still running at the end of the testing session, their times were recorded as 300 s. Three training sessions were performed one week prior to time off dox, and two test sessions were conducted weekly, with the final score being the highest time of the two test sessions. Experimenters were blinded to treatment group for hindlimb clasping and rotarod assessments.

### RNA extraction

RNA was extracted with Qiazol (Qiagen #79306) using the Qiagen RNeasy Mini Kit (Qiagen, #74104) and Precellys tissue homogeniser (Bertin Instruments, Montigny-le-Bretonneux, France). On-column DNase I digestion was conducted using RNase-free DNase I (Qiagen #79254). The concentration of extracted RNA was determined using a NanoRNA kit (Agilent #5067-1511) and Bioanalyzer (Agilent 2100, Santa Clara, CS, USA). cDNA was synthesized from 1 μg total RNA using the SuperScript™ VILO™ Master Mix (Thermo Fisher #11755050).

### RT2 PCR arrays and data analysis

For detection of differentially expressed genes of interest, cell stress and death RT^2^Profiler^TM^ PCR Arrays (SABioscience Corporation, #PAMM-003Z) were utilised. The SYBR-Green based RT-PCR arrays contain 84 genes plus 5 housekeeping genes and assay controls. cDNA was mixed with SYBR-green as per manufacturer instructions and loaded onto a 384 well plate, with each gene was assessed in quadruplicate. qPCR was performed on the Vii7 Real-Time PCR System (Applied Biosystems, Waltham, MA, *USA*). Fold-change analysis was performed using the ExpressionSuite Software (version 1.1), whereby each gene was determined by exponentiation of 2^-ΔΔCt^. A fold change of greater than 2 and p<0.05 was deemed significant. The ß2 microglobulin (B2m) gene was excluded from the housekeeping genes in analysis as the expression of B2m gene was significantly increased in the cortex of rNLS8 mice off Dox compared to controls (Supplementary Fig 1A), in line with previous findings in ALS mouse models *(60)*.

For hierarchical clustering, fold change data of genes of interest was processed using the web-based Morpheus software (Broad Institute, Cambridge, MA, USA). Classification of differentially expressed genes was performed to indicate co-regulated and functionally related genes. Gene ontology (GO) and Kyoto Encyclopedia of Genes and Genomes (KEGG) pathway enrichment analyses was performed using Metascape (version 3.0) with default parameters (minimum overlap = 3, p-value cutoff = 0.01, and minimum enrichment = 1.5) (61). Protein-protein interaction enrichment analysis was performed using the Molecular Complex Detection (MCODE) algorithm using Metascape to identify densely connected components from examined pathways with default parameters (62). Venn diagrams were constructed using InteractiVenn (63). Volcano plots were generated using Prism-GraphPad software (version 9).

### Real-time quantitative PCR (qPCR) assay

Target gene real-time qPCR analyses were conducted using SYBR Green Master Mix (Bioline #BIO-98005) with the LightCycler^®^ 480 System (Roche, Basel, Switzerland). The relative changes in gene expression were calculated based on the housekeeping gene beta-actin *(Actb)* according to the 2^-ΔΔCT^ method (64) and normalised to control or saline-treated control groups. The sequences of primers are available in Supplementary Table 1.

### Protein extraction and immunoblotting (IB)

Tissues were thawed on ice and then homogenised in 5× v/w RIPA lysis buffer (50 mM Tris, 150 mM NaCl, 1 % NP-40, 5 mM EDTA, 0.5 % sodium deoxycholate, and 0.1 % SDS, pH 8.0) containing 1 mM PMSF and protease and phosphatase inhibitor cocktails (Sigma, # 4906845001 and #11836170001) with three 1.4 mm Zirconium oxide beads (Bertin Instruments, #P000927-LYSK0-A) using Precellys tissue homogeniser (Bertin Instruments, #P000669-PR240-A). Samples were centrifuged at 4°C, 100,000 *g* for 30 min, and the supernatant was taken as the RIPA-soluble fraction. The remaining pellet was washed with RIPA buffer as above, this supernatant was discarded, and the resulting pellet was dissolved in 5× v/w urea buffer (7 M urea, 2 M thiourea, 4% CHAPS, and 30 mM Tris, pH 8.5) using the Precellys homogeniser and centrifuged at 22°C, 100,000 *g* for 30 min. This supernatant was taken as the RIPA-insoluble/urea-soluble fraction. Protein concentrations of the RIPA-soluble fractions were determined using the Pierce^TM^ BCA Protein Assay Kit (ThermoFisher Scientific #23225).

Protein samples of RIPA-soluble or RIPA-insoluble/urea-soluble fractions were separated by electrophoresis (120 V for 90 min) on a polyacrylamide gel containing 12% acrylamide in the presence of reducing agent (2-mercaptoethanol, Sigma #63689). After SDS-PAGE, proteins were transferred to nitrocellulose membranes (LI-COR Biosciences #P/N926-31092) and incubated in blocking solution (5% (w/v) BSA, 0.05% (w/v) Tween-20 in TBS (TBST)) and then incubated overnight in primary antibody diluted in the blocking solution. Primary antibodies used for immunoblotting were rabbit anti-TDP-43 polyclonal antibody (PAb) for detecting both human and mouse TDP-43 (Proteintech #10782-2-AP, 1:2000), rat anti-phospho-S409/410 TDP-43 monoclonal antibody (MAb) (Biolegend # 829901, 1:1000), mouse anti-CHOP Mab (Invitrogen #MA1-250, 1:1000), a rabbit anti-caspase-3 for detection of cleaved forms of caspase-3 (Abcam #9664, 1:1000), mouse anti-GAPDH Mab (Proteintech #60004-1-Ig, 1:10,000), and rabbit anti-GAPDH PAb (Proteintech #10494-1-AP, 1:2000). The nitrocellulose membranes were washed with TBST and incubated with IRDye secondary antibodies (LI-COR Biosciences, 1:20,000) for 1 hour. Protein bands were visualized using the Odyssey CLx Imaging System (LI-COR Biosciences, Lincoln, NE, USA) and quantified using Image Studio Lite software (LI-COR Biosciences, Lincoln, NE, USA). The relative changes of protein levels were calculated by quantifying the bands in RIPA soluble fractions relative to the band of the internal reference protein GAPDH, or by quantifying the bands in RIPA-insoluble/urea-soluble fractions relative to total protein and subsequently normalised to the control group or saline-treated control group.

### Histology and immunofluorescence labeling (IF)

Mice were perfused using PBS followed by 4% paraformaldehyde (PFA). The brain and lumbar spinal cord tissues were rapidly dissected and post-fixed in 4% PFA for 1 hour at room temperature and 4°C overnight. Post-fixed tissues were rinsed in phosphate-buffered saline (PBS) pH 7.4, dehydrated with a series of increasing concentrations of ethanol, and then embedded in paraffin. Embedded brains were sectioned at 10 μm thickness. For IF, paraffin-embedded sections were deparaffinised in xylene and rehydrated through a series of decreasing concentrations of ethanol. After rehydration, sections were subjected to antigen retrieval solution (10 mM sodium citrate, 0.05% Tween-20, pH 6.0) at 95°C for 10 min and then were allowed to cool. Sections were then blocked in 1% BSA in PBS containing 0.25% (v/v) Triton X-100 (PBST) and incubated in primary antibodies overnight at 4°C. Primary antibodies used for IF were rabbit anti-TDP-43 polyclonal antibody (PAb) for detecting pan-TDP-43 for both human and mouse TDP-43 (Proteintech #10782-2-AP, 1:1000), and rat anti-glial fibrillary acidic protein (GFAP) Mab (ThermoFisher Scientific #13-0300, 1:500). Sections were then washed in PBST and incubated with fluorophore-conjugated secondary antibodies, including donkey anti-rat conjugated Alexa-488 (ThermoFisher Scientific #21208, 1:1000) and donkey anti-rabbit conjugated Alexa-647 (ThermoFisher Scientific #48272, 1:1000). Nuclei were stained using 4’,6-diamidino-2-phenylindole (DAPI, Sigma-Aldrich #D8417, 1:1000).

### Fluorescence Microscopy and Image analysis

Images for IF sections were acquired using an epifluorescence microscope Axio Imager (Zeiss) under 20 x objective (0.8 NA / 0.55 mm WD), providing a pixel size of 0.323 μm using Zen imaging software (Zeiss). Images were background-corrected using ImageJ software with the rolling ball background subtraction algorithm and were then quantified for protein levels of TDP-43 using CellProfiler software (Broad Institute, version 4.2.1) as previously published (65).

### Statistical analysis

For two-value data, statistical analyses were conducted using a two-tailed t-test. One-way ANOVA was used for analyses of datasets with three values with Bonferroni’s post hoc test using Prism-GraphPad software (version 9). Statistical significance is indicated as \**p* < 0.05, \*\**p* < 0.01, and \*\*\**p* < 0.001. All data are presented as mean ± standard error of the mean (SEM).

## Supplementary Material

**Supplementary Fig. 1.**
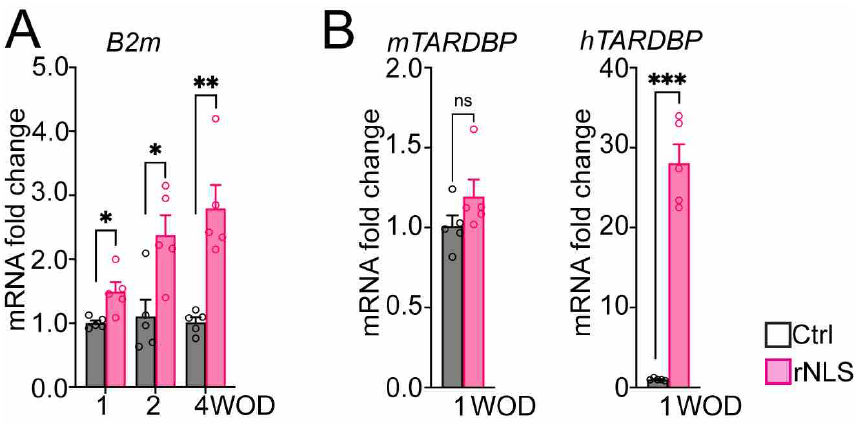
Real-time qPCR assessed the expression of genes in the cortex of rNLS8 mice at 1, 2 or 4 weeks off Dox (WOD). **A**. B2m. **B**. Mouse (m) and human (h) TARDBP. Actb was used as the housekeeping gene for normalisation. *n = 5. Mean + SEM. *p < 0.05, **p < 0.01, and ***p < 0.001.*

**Supplementary Fig. 2.**
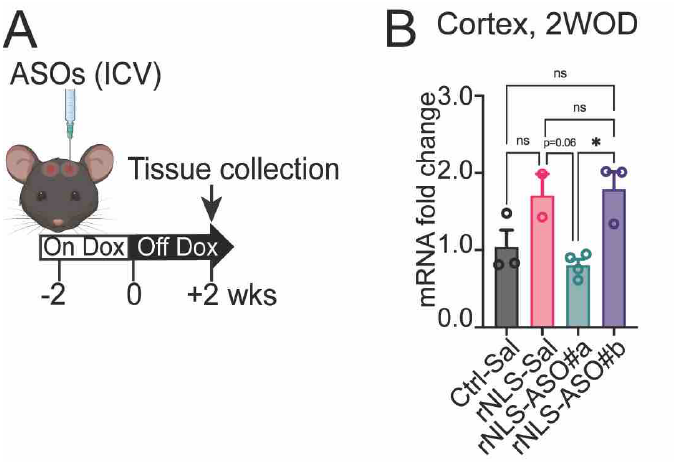
Chop ASO#a inhibits Chop gene expression in rNLS8 mice at 2 weeks off Dox (WOD). **A**. Experimental schema. B. Real-time qPCR results demonstrate knockdown of Chop gene by ASO#a (previously reported as ASO#3, see Table 2) in the cortex of treated rNLS8 mice, but not by ASO#b (previously reported as ASO#5, see Table 2). Saline-treated control mice (n = 3), saline-treated rNLS8 mice (n = 2), ASO#a-treated rNLS8 mice (n = 4) and ASO#b-treated rNLS8 mice (n = 3). Chop ASO#a treatment prevented increases of CHOP mRNA in rNLS8 mice, as previously reported in other models. Data are normalised to Actb as the housekeeping gene. Sequences of ASOs are available in Table 2. Mean + SEM. * indicates p <0.05 (one-way ANOVA with Bonferroni’s post hoc test).

**Supplementary Table 1.**
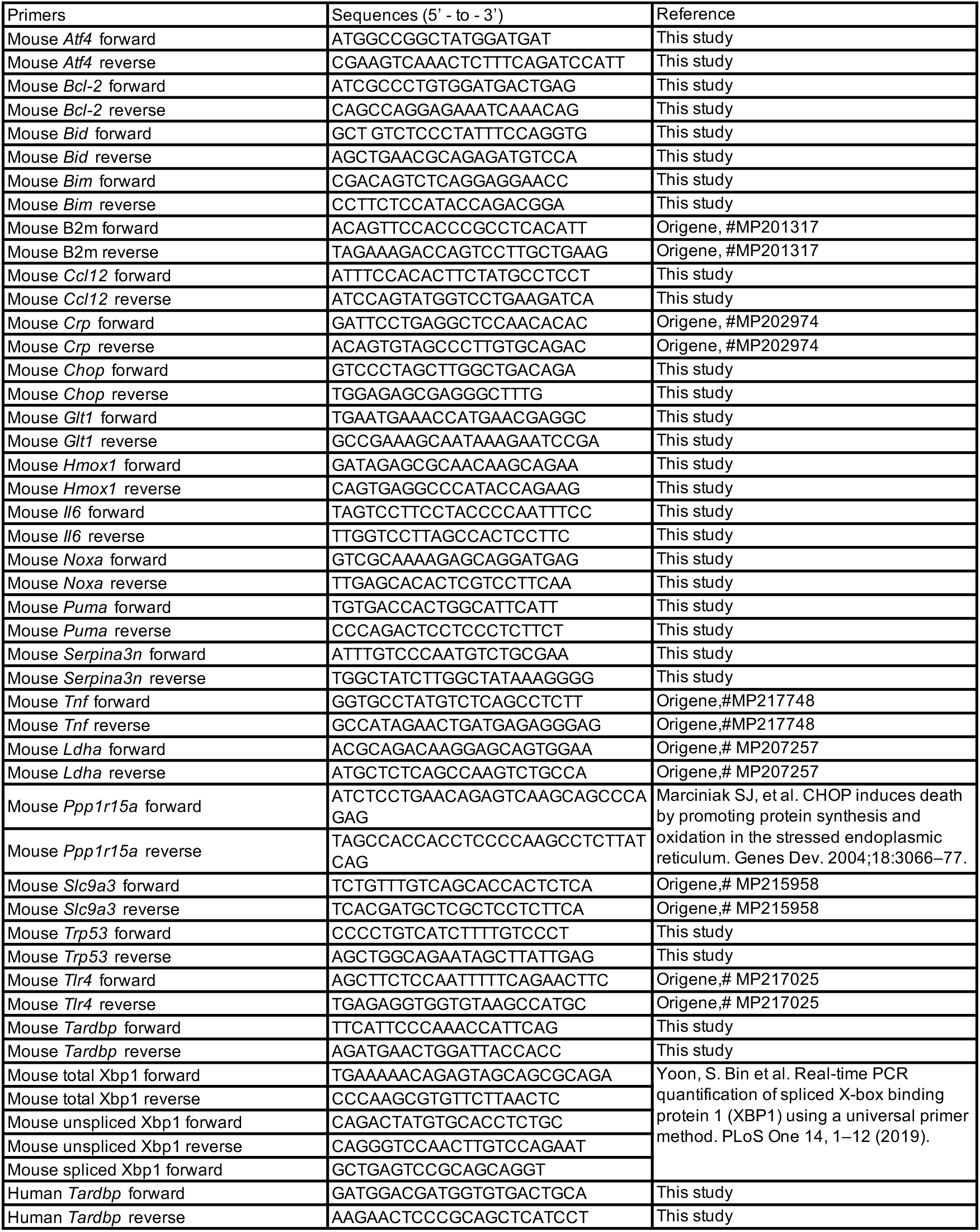
Sequences of primers used for real-time qPCR

**Supplementary Table 2.**
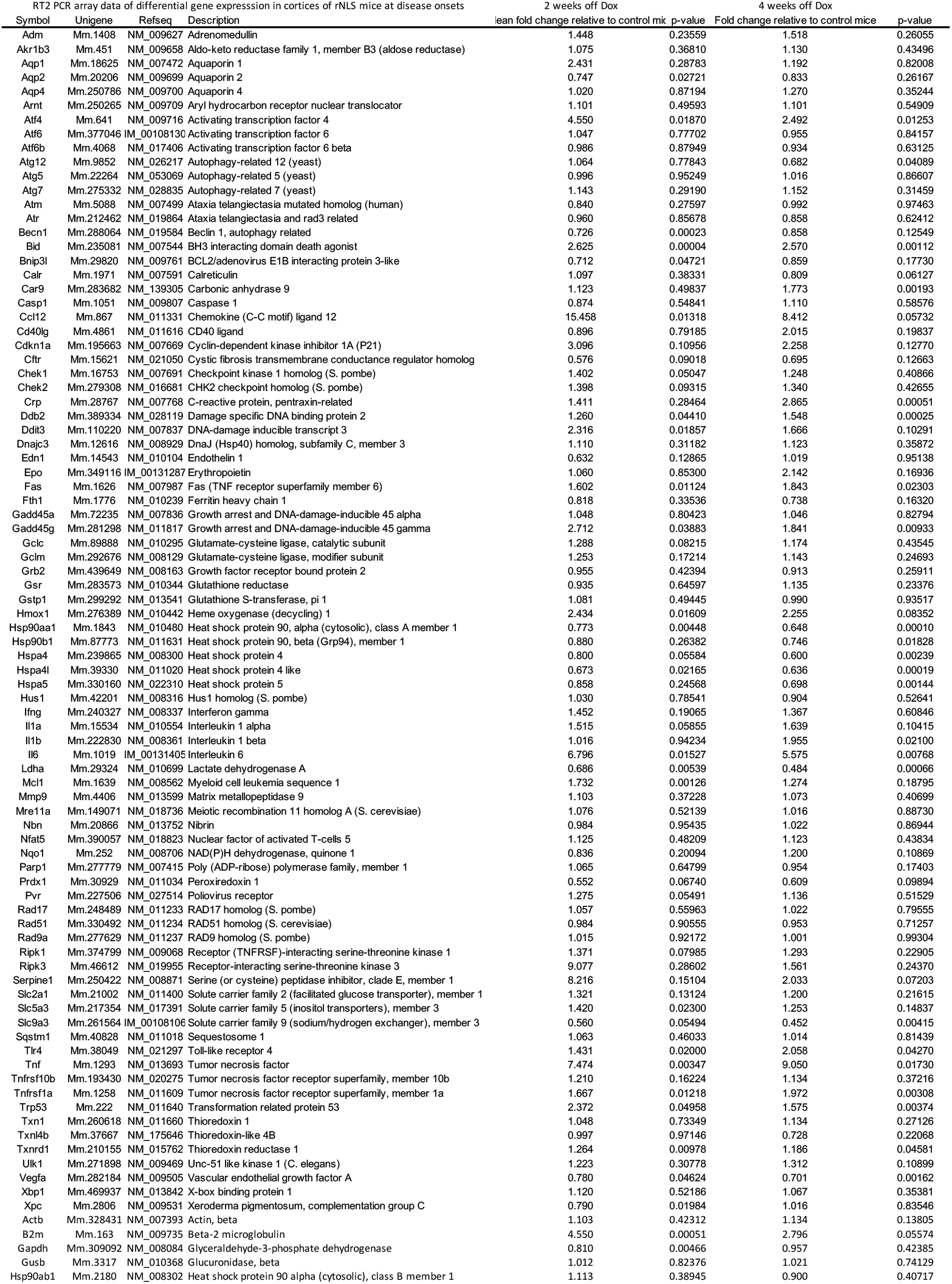
Raw RT^2^ PCR array data of differential gene expression in cortex of rNLS8 mice at 2 weeks and 4 weeks off Dox

## Acknowledgement

The authors thank the staff of animal and behavioural facilities at Queensland Brain Institute and Macquarie University for animal husbandry and surgery support, and the QBI Histology Facility and the QBI Advanced Microscopy Facility for their support and assistance in this work. Figures were constructed using Biorender.com.

## Ethical Approval

All animal procedures were conducted with approved from the Animal Ethics Committee of Macquarie University (#2016-026), and the Animal Ethics Committee of The University of Queensland (#QBI/131/18), in accordance with the Australian Code of Practice for the Care and Use of Animals for Scientific Purposes.

## Author contributions

WL, ALW, SL, FR, PJN, KL and AKW designed research; WL, ALW, HBW, LMSM and AKW performed research; WL, ALW, HBW, SL, RSG and AKW contributed unpublished reagents/analytic tools; WL, ALW, HBW, RSG and AKW analysed data; WL, ALW, RSG and AKW wrote the paper. All authors read and approved the final manuscript.

